# Evidence for a transfer-to-trap mechanism of fluorophore concentration quenching in lipid bilayers

**DOI:** 10.1101/2024.02.16.580699

**Authors:** Sophie A. Meredith, Yuka Kusunoki, Stephen D. Evans, Kenichi Morigaki, Simon D. Connell, Peter G. Adams

## Abstract

It is important to understand the behaviours of fluorescent molecules because, firstly, they are often utilized as probes in biophysical experiments and, secondly, they are crucial cofactors in biological processes such as photosynthesis. A phenomenon called ‘fluorescence quenching’ occurs when fluorophores are present at high concentrations but the mechanisms for quenching are debated. Here, we used a technique called ‘in-membrane electrophoresis’ to generate concentration gradients of fluorophores within a supported lipid bilayer (SLB), across which quenching was expected to occur. Fluorescence lifetime imaging microscopy (FLIM) provides images where the fluorescence intensity in each pixel is correlated to fluorescence lifetime: the intensity provides information about the location and concentration of fluorophores and the lifetime reveals the occurrence of energy-dissipative processes. FLIM was used to compare the quenching behaviour of three commonly-used fluorophores: Texas Red (TR), nitrobenzoaxadiazole (NBD) and 4,4-difluoro-4-bora-3a,4a-diaza-*s*-indacene (BODIPY). FLIM images provided evidence of quenching in regions where the fluorophores accumulated but the degree of quenching varied between the different fluorophores. The relationship between quenching and concentration was quantified and the ‘critical radius for trap formation’, representing the relative quenching strength, was calculated as 2.70, 2.02 and 1.14 nm, for BODIPY, TR and NBD, respectively. The experimental data supports the theory that quenching takes place via a ‘transfer-to-trap’ mechanism which proposes, firstly, that excitation energy is transferred between fluorophores and may reach a ‘trap site’ resulting in immediate energy dissipation and, secondly, that trap sites are formed in a concentration-dependent manner. Some previous work suggested that quenching occurs only when fluorophores aggregate, or form long-lived dimers, but our data and this theory argues that traps may be ‘statistical pairs’ of fluorophores that exist only transiently. Our findings should inspire future work to assess whether these traps can be charge-transfer states, excited state dimers or something else.

## INTRODUCTION

Fluorescence is a fundamental physical process that occurs after a molecule absorbs light, enters a higher-energy excited state and then releases that energy as photon emission (1). A great variety of fluorescent molecules (fluorophores) exist, ranging from synthetic organic compounds (2) to natural biomolecules (3), and are often used as a labelling tool in biophysics research. Despite the wide-ranging use of fluorophores their behaviour is not fully understood, particularly the process of ‘quenching’ that can occur when multiple fluorophores interact together (4). Concentration-dependent self-quenching is a phenomenon observed in experiments where the fluorescence intensity measured per-fluorophore decreases non-linearly as the concentration of fluorophores increases (5-10). In biophysical measurements, unexpected quenching can be a problem if it disrupts fluorescence assays (11-13). In the biological process of photosynthesis, the quenching of excited states plays an important role within light-harvesting (LH) pigment-protein complexes (14, 15). These LH complexes perform a balancing act: they contain very high concentration of chlorophyll pigments (∼250 mM) but, yet, they avoid energy loss and quenching under normal conditions (14). Interestingly, certain LH proteins switch from an active state to a highly quenched state when the biological organism is exposed to high-intensity sunlight, as a means to safely dissipate excess energy, and there is debate about the protein structural changes and photophysical pathways involved (16-25). Based on the need to understand both synthetic fluorophores and natural LH proteins (26-28), it is of fundamental importance to understand the physics behind fluorescence quenching.

Biological LH proteins have a relatively high level of complexity, containing tens of pigments of a variety of types within a relatively large system, and this can make modelling energetic transitions challenging (18, 29, 30). Therefore, it is instructive to perform measurements on simpler model systems where there is a single type of fluorophore in a random distribution. Previous experimental studies have assessed the extent of quenching occurring for model systems such as pigments dissolved in organic solvents (a 3-D system) and pigments within a lipid membrane (a 2-D system). For example, investigations of chlorophylls (5, 6) and synthetic organic fluorophores (8-10) found that the onset of quenching typically occurs when the concentration of pigments exceeds ∼5 mM leading to average inter-pigment distances <5 nm. Previous theoretical studies have attempted to test the structural basis of concentration-based quenching by considering networks of fluorophores, developing mathematical models (9, 31-35) and performing computational simulations (34, 36-39). Based on the findings of both experiment and theory, there are two types of energy ‘traps’ proposed to cause quenching in simple systems involving a single type of fluorophore: (i) ground-state dimers of fluorophores that are relatively long-lived due to strong chemical or physical attractions (10, 40-42), (ii) ‘statistical pairs’ of fluorophores that do not have any attractive interactions in their ground states and only interact transiently when one of the pair of fluorophores enters an excited state (7, 32, 43). It is logical that ground-state aggregates of a fluorophore would lead to quenching because direct inter-molecular contact could lead to efficient energy dissipation via electron transfer processes (44) but such aggregation would require strong attractive interactions, which seem unlikely to occur for all types of fluorophores in random distributions. In order to better understand the behaviour of fluorophores, it is important to determine which of these quenching mechanisms is the best representation in each case.

For experimental investigations of fluorescence quenching, a controllable system of fluorophores is required where the inter-fluorophore distances can be varied. The results can then be more accurately compared to the theoretical models (27). In past studies, a series of fluorophore samples across a concentration range would be prepared, for example 0.01-10 mM chlorophyll in organic solvents (5), and analysed with fluorescence spectroscopy. Each individual sample resulted in a single datapoint on a quenching-versus-concentration graph so it was time-consuming to assess complex trends. We recently published a new method for preparing and analysing a range of fluorophore concentrations within a single sample to allow for a deeper study of fluorescence quenching (45). In this work, the commonly used fluorophore Texas Red (TR) was incorporated within supported lipid bilayers (SLBs) at a relatively low concentration and then an electric field was applied in order to generate a concentration gradient of TR. The novelty here was, firstly, to use Fluorescence Lifetime Imaging Microscopy (FLIM) to take images where the fluorescence lifetime was correlated to the fluorescence intensity of TR, allowing quenching to be quantified (45), and secondly, to exploit the ‘in-membrane electrophoresis’ technique (46-51) to study quenching. We found that electrophoresis generated a roughly exponential gradient of TR concentration across a SLB and that concentrations of >2% TR (mol/mol% relative to total lipid) were correlated to significant quenching (45). A knowledge gap highlighted by this work was that the molecular basis for the energy dissipation underlying quenching was unknown. Furthermore, only one type of fluorophore was studied and we noted that different types of fluorophores could have different quenching behaviour due to different chemical structures and physical characteristics.

In this paper, we use in-membrane electrophoresis and FLIM to compare three different organic fluorophores, allowing us to contrast the quenching behaviour of different fluorophore chemistries. To assess the mechanism of quenching, our experimental results are compared with a theoretical model for ‘transfer-to-trap’ quenching.

## MATERIALS AND METHODS

### Theory

#### Theoretical model for fluorophore quenching as a function of concentration in 2-D

At its simplest level, our experimental model system produces a defined quantity of fluorophores in a random arrangement within a 2-D plane. One benefit of localizing fluorophores to a very thin film is that reabsorption (inner filter) effects, that distort the spectra of concentrated 3-D solutions, are avoided (35). This model system is a self-assembled 2-D film of lipids containing certain quantities of fluorescent probes (**Fig. 1 *A***), but the same theory will apply to other 2-D systems. For simplicity, we consider the energy transfer within only one leaflet of the lipid bilayer and assume that both leaflets have the same distribution of fluorophores. To identify the molecular mechanism of fluorophore self-quenching, we apply mathematical models that were previously suggested to represent quenching effects and adapt them for our experimental system. The majority of equations have been derived previously (as cited), here they are applied to predict quenching relationships for our fluorophores-of-interest and to derive an expression for analysis of FLIM data (Eq. 22).

**Figure 1.**
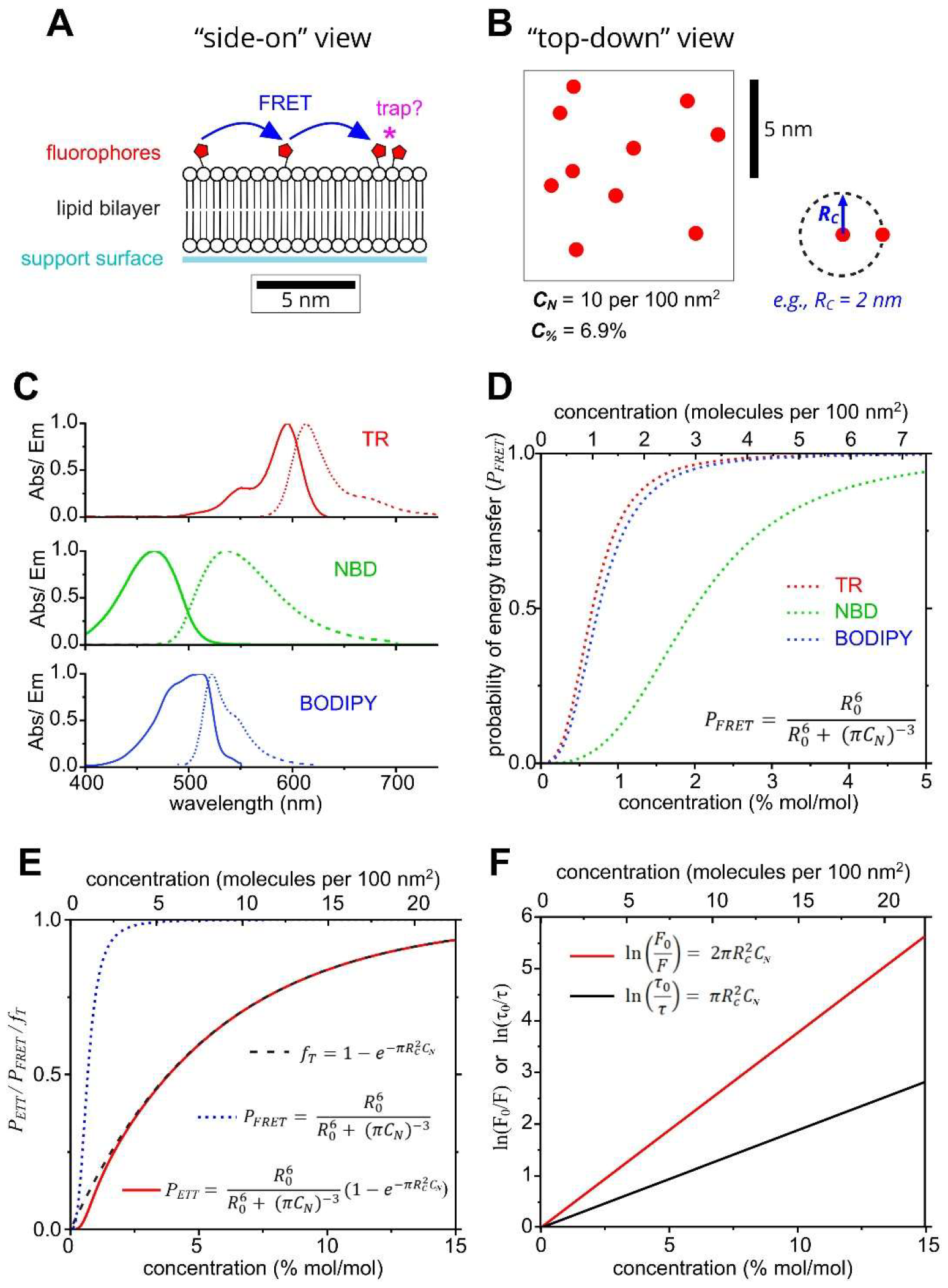
Schematics and calculations of energy transfer and trapping properties for TR, NBD and BODIPY based on theoretical models. **(A)** Cartoon of the model membrane system containing a small number of fluorophores tethered to lipid headgroups, drawn to scale (viewed from a ‘side-on’ perspective). A hypothetical trap site formed by a ‘statistical pair’ is denoted by a pink asterisk. **(B)** A top-down view of a 10 × 10 nm box containing 10 fluorophores, at the same scale as (A). This represents a number density of C_N_ = 0.1 nm^-2^, equivalent to C_%_ = 6.9% mol/mol. The fluorophore is assumed to occupy the same area as a lipid (0.69 nm^2^). **(C)** Absorption and fluorescence spectra of TR, NBD and BODIPY (as labelled) used to calculate the Förster radius of each fluorophore. **(D)** Theoretical plots of the probability of FRET between two fluorophores as a function of their concentration, according to the equation shown. **(E)** Theoretical plots of P_FRET_, f_T_ and P_ETT_ for TR using R_0_ = 5.71 nm and R_C_ = 2.00 nm (as labelled). A schematic depicting R_C_ = 2 nm is shown in (B). **(F)** Theoretical plots of the logarithm of the relative fluorescence intensity or lifetime as a function of fluorophore concentration, as expected for quenching due to the transfer-to-trap model.

First, the distances at which interactions between fluorophores could take place were considered. Previous microscopy measurements of lipid bilayers containing a range of fluorophore concentrations have shown that significant fluorescence quenching occurs at a concentration of 1% fluorophores, where % represents the percentage of total lipid molecules that are fluorescently tagged (45). The fluorophore concentration can be converted from this mole-to-mole percentage, as typically reported in experiments, to a number density because this is more useful for spatial considerations:

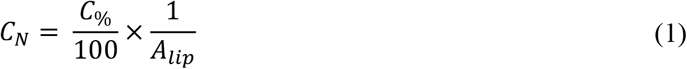

where *C*_*N*_ and *C*_*%*_ are the concentration of fluorophores in molecules per nm^2^ (i.e., number density) and in mol/mol %, respectively, and *A*_*lip*_ is the area occupied by a single lipid (estimated to be 0.69 nm^2^ (52)). The cartoon in **Fig. 1 *B*** shows an example *C*_*N*_ of 0.1 per nm^2^ equivalent to *C*_*%*_ ∼6.9%.

Next, the average distance between molecules (*r*, centre-to-centre), can be related to their concentration via the area of a circle drawn around the molecule:

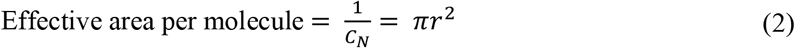

Thus, for a membrane containing a typical concentration of 1% mol/mol fluorescent lipids, the average distance between fluorophores can be calculated as 4.69 nm.

Now that concentrations and distances have been considered, a theoretical model for concentration quenching can be applied. One possibility is that an excited fluorophore may diffuse into direct contact with a nearby fluorophore whereby the fluorescence is quenched by the collision of the molecules, sometimes termed ‘collisional quenching’. To test this possibility, the properties of the TR fluorophore were used to calculate the mean displacement due to diffusion, *<x>*, over a time equal to the fluorescence lifetime, as a representation of the period that an excited state would typically persist:

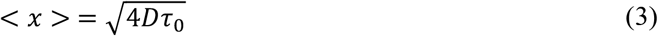

where *D* is the diffusion constant (*D* = 2.27 μm^2^/s, **Fig. S1**) and *τ*_*0*_ is the fluorescence lifetime for TR (*τ*_*0*_ = 4.23 ns, **Table S3**). *<x>* was found to be ∼0.20 nm, significantly lower than the fluorophore separation distance of 4.69 nm, clearly suggesting that molecular collisions would occur infrequently in this system, so cannot explain the significant quenching (45). Thus, we must consider models that involve the transfer of excited states over a distance of several nanometres. The most likely means is the non-radiative transfer of energy via resonance interactions as described by Förster theory (53), whereby an individual excited state can ‘hop’ between fluorophores.

Numerous studies have proposed that self-quenching processes involve a combination of two important processes: excited state transfer and energy dissipation. The overarching idea is that an exciton quasiparticle may rapidly migrate between multiple fluorophores and finally become quenched at a trap site (**Fig. 1 *A***) (9, 32). This will affect the observed fluorescence intensity and lifetime, as follows:

(i) Two fluorophores that are closer than a critical distance will form a trap site. Traps are non-fluorescent by definition (either photon absorption is forbidden or non-radiative dissipation of energy is immediate). Thus, the overall fluorescence intensity is reduced by the fraction of fluorophores involved in trap sites, as follows:

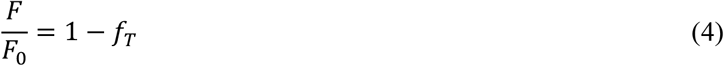

where *F*_*0*_ is the original fluorescence intensity, *F* is the fluorescence intensity after this type of quenching and *f*_*T*_ is the fraction of fluorophores involved in traps. This is sometimes termed ‘static quenching’.
(ii) In addition, fluorophores that are not part of traps may become excited and then transfer the exciton to other fluorophores via Förster Resonance Energy Transfer (FRET). If FRET causes an exciton to reach one of the trap sites described in (i) then the energy is immediately dissipated. In this process, the transfer mechanism provides a route for quenching to occur over relatively long distances (several nanometres) and the trap provides a process for energy dissipation. This dissipative pathway is an alternative to fluorescence, resulting in a reduction in both the fluorescence intensity and fluorescence lifetime. This is termed ‘transfer-to-trap quenching’. The relative change in lifetime should be equal to:

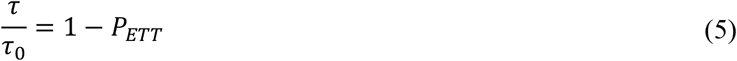

where τ_O_ is the original fluorescence lifetime, τ is the fluorescence lifetime after quenching and *P*_ETT_ is defined as the probability of excitation transfer to a trap site.

The fluorescence intensity will also be reduced in the same manner:

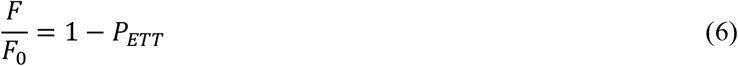

These two processes will occur simultaneously in a system that undergoes concentration-dependent quenching, so that the fluorescence intensity is reduced by (i) and (ii), whereas the lifetime is only affected by (ii). So, the overall change in fluorescence intensity will be represented by the combination both processes and the complete expression is the combination of Eq. 4 and Eq. 6:

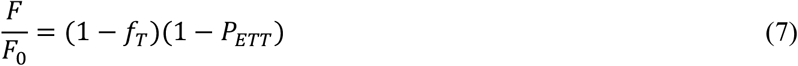

Now, the physical origin of *f*_T_ and *P*_ETT_ must be considered. Our theoretical model will consider a random distribution of molecules that represent the situation of traps being ‘statistical pairs’ of fluorophores. If this minimal model is sufficient to explain experimental data on fluorescence quenching then it would imply that concentration quenching does not require molecular aggregation. For a random distribution of particles in a 2D-plane with a concentration, *C*, the number of particles separated by a distance less than *R* can be derived statistically (32, 54). In a quenching model system, this approach can be used to determine the fraction of fluorophores that are part of traps, *f*_*T*_, as:

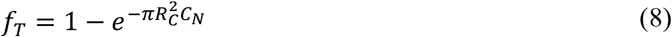

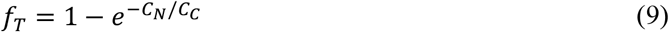

where *R*_*C*_ is a property of the fluorophore known as the ‘critical radius for trap formation’ that represents the propensity of two fluorophores to form a trap site when in close proximity (7, 32). *R*_*C*_ is defined as the distance at which *f*_T_ = (*1 – e*^*–1*^) ≈ 0.63, and this leads to an equivalent ‘critical concentration’, *C*_C_ (i.e., at *C*_N_ = *C*_C_ the *f*_T_ ≈ 0.63). Therefore, a fluorophore that has a higher *R*_*C*_ has the ability to form traps more readily and will self-quench more strongly. These traps can be considered as quasi-stable and immobile for the duration of an excited state due to the slow lateral (<x> ∼ 0.2 nm) and rotational diffusion of fluorophores compared to their fluorescence lifetime.

The probability that any given exciton reaches a trap site can then be expressed as

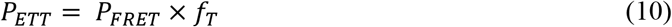

where *P*_FRET_ is the probability of resonance energy transfer, as described by Förster theory (53), and *f*_*T*_ is the fraction of fluorophores that exist as traps as defined earlier. This expression assumes that an excited molecule only undergoes FRET to its nearest-neighbour, justified by multiple studies (33, 36). To assess whether transfer-to-trap quenching could be a good model for our system, graphs of the theoretical *P*_FRET_, *f*_T_ and *P*_ETT_ were calculated as a function of the concentration of fluorophores that can be studied experimentally, as described below.

*P*_FRET_ depends upon the strength of dipole-dipole coupling between a potential donor and acceptor molecular and this has a strong distance-dependence. The potential for dipole-dipole coupling (and also energetic coupling) can be encapsulated in the expression for Förster radius, *R*_*0*_ (53):

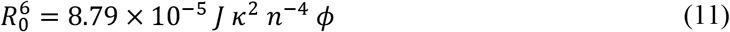

where *κ* represents the relative orientation of the donor and acceptor transition dipoles, *n* is the optical refractive index of the medium, *ϕ* is the fluorescence quantum yield of the donor, and *J* is the spectral overlap integral between the donor emission and acceptor absorption. In other words, *R*_*0*_quantifies the potential of a donor-acceptor pair to transfer excitation energy where a higher value leads to greater FRET. *R*_*0*_ is defined as the inter-fluorophore separation distance at which *P*_FRET_ = 50%.

*R*_*0*_ was calculated for three different fluorescent probes that were used in our experiments with lipid bilayers: Texas Red (TR), nitrobenzoaxadiazole (NBD) and 4,4-difluoro-4-bora-3a,4a-diaza-*s*-indacene (BODIPY). In our scenario where the exciton donor and acceptor are identical molecules, a value for *J* can be calculated as the area overlap between the measured emission and absorption spectra for each fluorophore (9, 10), as shown in **Fig. 1 *C***. Graphical integration of these overlaps found values of 3.08, 0.28 and 2.13 × 10^15^ M^-1^cm^-1^nm^4^ for TR, NBD and BODIPY, respectively. For our system, the fluorescent moiety was assumed to be randomly-orientated due to the effects of both lateral diffusion of the lipids and rotational diffusion of the tethered fluorophores, equating to a value for *κ*^*2*^ of 2/3 (55). The refractive index of the optical medium was given a value of 1.45, halfway between that of water (1.33) and lipids (1.55), as in previous estimates (56). The fluorescence quantum yield of TR, NBD and BODIPY has been previously determined as 0.93, 0.40, 0.99 (9, 57, 58). Using these values and Eq. 11, the Förster radius *R*_*0*_ was found to be 5.73, 3.32 and 5.41 nm for TR, NBD and BODIPY, respectively. Therefore, the potential for FRET is roughly similar for TR and BODIPY and lower for NBD.

Next, *P*_FRET_ was calculated as a function of concentration for the three fluorophores using the values determined for *R*_*0*_ and a range of concentrations, *C*_*N*_, and the following expression from Förster theory:

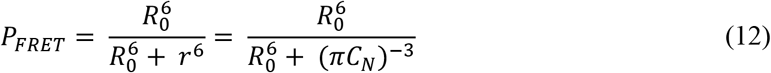

The results of these calculations are plotted in **Fig. 1 *D***, where it can be seen that for all fluorophores the probability of FRET increases rapidly as a function of concentration before approaching unity at *C*_*%*_ ∼2% for TR and BODIPY and ∼5% for NBD. This result shows that, for the typical concentrations of fluorophores found in lipid bilayers, excitation energy transfers are highly likely to occur, leading to efficient random migration of excitons throughout the membrane.

Now, starting from the definition in Eq. 10, *P*_*ETT*_ can be re-written by combining Eq. 12 and Eq. 8, as:

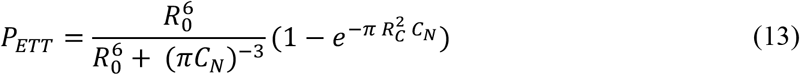

It is important to note that this theoretical quenching probability (*P*_ETT_) is equivalent to the experimentally observable quenching behaviour:

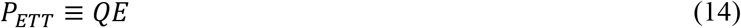

where *QE* is the ‘quenching efficiency’ calculated from experimental measurements of fluorescence lifetime (i.e., spectroscopy), as follows:

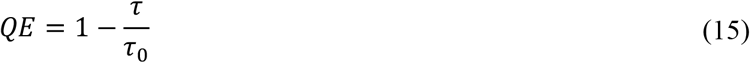

To generate graphs of the theoretical quenching behaviour, the values for *f*_*T*_, *P*_*FRET*_and *P*_*ETT*_ (Eq. 8, 12 and 13, respectively) were calculated for a range of TR concentrations. These theoretical calculations used the value for *R*_*0*_ (TR-TR) = 5.73 nm (as determined above) and a placeholder value for critical radius of trap formation, *R*_*C*_ = 2.0 nm, and inputted a range of *C*_%_= 0-15%. This value for *R*_*C*_ is a reasonable first estimate, based on previous reports on other fluorophores (9, 10), but this parameter will be quantified experimentally later in this report. **Fig. 1 *E*** shows the theoretical *f*_*T*_ (*black*), *P*_*FRET*_ (*blue*) and *P*_*ETT*_ (*red*) versus concentration curves for TR. All three curves tend asymptotically towards unity, but that the rate of this approach varies drastically. *P*_*FRET*_ increases steeply with concentration before asymptotically tending towards unity at concentrations above ∼2%, whereas *f*_*T*_ and *P*_*ETT*_ increase with shallower gradients and only begin to tend towards unity at much higher concentrations, over 15%. Most significantly, the theoretical plots demonstrate the similarity between the *P*_*ETT*_ and *f*_*T*_ curves, which are almost perfectly overlaid for the concentrations of 1.5 to 15%. At lower concentrations, *P*_*ETT*_ deviates from *f*_*T*_ and at ∼1% TR there is a difference of ∼25%. This suggests that transfer-to-trap quenching should be dominated by the number of traps in the membrane for TR and that at sufficiently high fluorophore concentrations of >1.5% TR it is reasonable to assume that *P*_*ETT*_ *≈ f*_*T*_ (i.e., that *P*_*FRET*_ ≈ 1). The multiple redundant pathways for FRET between fluorophores towards a trap site increase the overall efficiency and make this assumption even more reasonable (i.e., multiple options for exciton hopping where several fluorophores are nearby). Furthermore, energy transfer could occur between fluorophores in different leaflets of the lipid bilayer, given a membrane width of ∼4 nm, and this would further increase *P*_*FRET*_.

Having established the potential of these theoretical models, we wished to apply them to our experimental observations of fluorescence, in order to gain a better understanding of the quenching mechanism. Using the approximation that *P*_*ETT*_ *= f*_*T*_, simplified relationships can be derived that relate a reduction in fluorescence lifetime and intensity to these statistical models for quenching (9).

Firstly, considering that the fluorescence lifetime is affected by transfer-to-trap quenching (and not static quenching), starting with Eq. 5, and substituting in *P*_*ETT*_ *= f*_*T*_ and Eq. 8, the relative change in lifetime is:

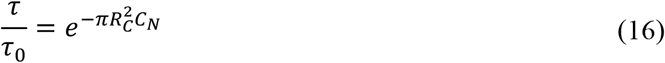

This can be written in a semi-logarithmic format to provide a linear relationship between fluorophore concentration and the amount of quenching:

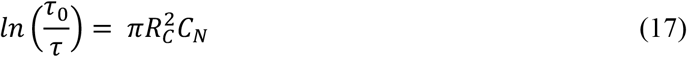

or, alternatively, the change in lifetime can be related to quenching efficiency (combining Eq. 15-16):

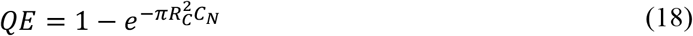

Secondly, considering that the fluorescence intensity is affected by both transfer-to-trap quenching and static quenching as in Eq. 7 and then simplifying with the *P*_*ETT*_ *= f*_*T*_ assumption:

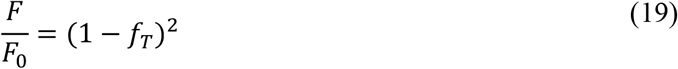

Therefore (using Eq. 8):

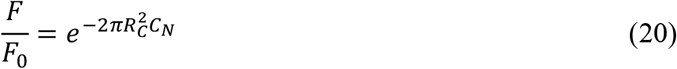

and, again, by rearranging and taking the natural logarithm:

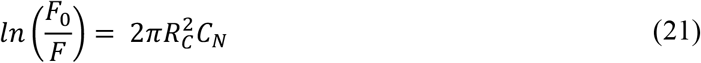

Plots of *ln(τ*_*0*_*/τ)* (Eq. 17) and *ln(F*_*0*_*/F)* (Eq. 21) as a function of concentration, across a typical range, are shown in **Fig. 1 *F***. The linear relationships, expected for fluorescence quenching, will later be compared to experimental results of in-membrane electrophoresis, to determine whether transfer-to-trap quenching occurs for lipid bilayers containing either TR, NBD or BODIPY. Furthermore, the gradient determined from a plot of *ln(τ*_*0*_*/τ)* vs. *C* will be used to determine the critical radius for trap formation, *R*_*C*_, to quantify the quenching strength of all three fluorophores.

Finally, we can derive an equation for the expected ‘non-quenched’ fluorescence intensity (*F*_*0*_) in terms of the observed fluorescence intensity (*F*) and the fluorescence lifetime ratio *τ/τ*_*0*_ by combining Eq. 17 and Eq. 21 and rearranging for *F*_*0*_:

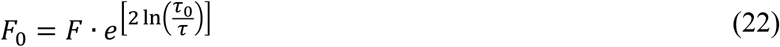

The derivations and calculations in this section show how the mechanism of transfer-to-trap quenching can be theoretically modelled for the three fluorophores studied and provide expressions for estimating the ‘critical radius of trap formation’. In the following sections, these mathematical models will be applied to experimental measurements of fluorescence intensity and lifetimes, to judge the quenching strength and mechanism for each fluorophore.

### Experimental Methods

#### Preparation of membrane corrals

1,2-bis(10,12-tricosadiynoyl)-*sn*-glycero-3-phosphocholine (Diyne-PC) lipids and 1,2-dioleoyl-*sn*-glycero-3-phosphocholine (DOPC) lipids were purchased as solids from Avanti Polar Lipids. The fluorescently-tagged lipids Texas Red 1,2-dihexadecanoyl-*sn*-glycero-3-phosphoethanolamine (TR-DHPE), 1,2-dipalmitoyl-*sn*-glycero-3-phosphoethanolamine-*N*-(7-nitro-2-1,3-benzoxadiazol-4-yl) (NBD-DHPE) and *N*-(4,4-difluoro-5,7-dimethyl-4-bora-3a,4a-diaza-*s*-indacene-3-propionyl)-1,2-dihexadecanoyl-*sn*-glycero-3-phosphoethanolamine (BODIPY FL DHPE) were purchased as solids from Invitrogen (Thermo Fischer Scientific), Avanti Polar Lipids, and Invitrogen, respectively. Template patterns were prepared as described in previous publications (45, 59, 60). Briefly, SLBs of Diyne-PC were formed on glass substrates by vesicle spreading and then polymerization was conducted by UV irradiation through a photomask of the desired pattern, here, a 2-D array of 100 × 100 μm boxes. Non-polymerized Diyne-PC molecules were removed with a detergent solution (0.1 M SDS), forming empty corrals surrounded by polymerized lipid bilayers. Lipid vesicles comprised of the specified ratio of fluorescent lipids to DOPC lipids were generated with standard probe sonication procedures in pure water. To form membrane corrals, a suspension of lipid vesicles (at a concentration of ∼0.5 mg/mL total lipid) was incubated with a template pattern for 20 min and then rinsed with a low ionic strength buffer (purified water, adjusted to pH 7.5 using <0.1 mM HCl).

#### In-membrane electrophoresis

A custom-built electrophoresis chamber (50, 51) was used to hold a glass substrate under aqueous buffer in a suitable position for microscopy and allow the application of a controlled E-field. The chamber was connected to a peristaltic pump via liquid outlets and a continuous 0.25 mL/min flow of buffer was provided during electrophoresis experiments to prevent the build-up of bubbles at the electrodes. Electrophoresis was performed on patterned membranes by applying an E-field (in-plane with the SLB) of 45 V/cm and monitored using a voltmeter throughout all experiments.

#### Fluorescence Lifetime Imaging Microscopy (FLIM)

FLIM was performed using a Microtime 200 time-resolved fluorescence microscope (PicoQuant GmbH). This system used an Olympus IX73 inverted optical microscope as a sample holder with light passing into and exiting various filter units for laser scanning, emission detection, and timing electronics. An excitation laser (561 or 485 nm) was driven in pulsed mode by a PDL 828 Sepia II burst generator module at a repetition rate of 10 MHz (pulse width 70-100 ps). An appropriate dichroic mirror and bandpass emission filter was used to select the detection range. For the TR fluorophore the excitation was at 561 nm and the collection between 590-650 nm, whereas for NBD and BODIPY the excitation was at 485 nm and the collection between 500-540 nm. The detector was a hybrid Photomultiplier Tube and the instrument response function was measured to have full-width at half-maximum of 100-120 ps. An excitation fluence of 0.012 mJ cm^−2^ was used for all measurements, which produced sufficient fluorescence signal while limiting any singlet-singlet annihilation events (see **Fig. S1**). Images were acquired by scanning the laser using a galvanometric (FLIMbee) scanner and accumulating many frames of the same region (1 frame = 3.2 s). A standard FLIM image was 25 frames (80 s of exposure), an optimal acquisition time that balanced the need for obtaining a strong fluorescence signal with the requirement to minimize photobleaching (45). The possibility of oxygen-dependent redox effects that may affect fluorophore photophysical properties was minimized by de-gassing all buffer solutions prior to use. Initial analysis of FLIM data was performed with SymPhoTime software (PicoQuant). The mean amplitude-weighted lifetime of images or pixels, <τ>, was calculated by generating fluorescence decay curves from accumulated photons and modelling the curve as a multiexponential decay function (excellent fits were achieved for all data, with chi-squared values <1.1 and low residuals). Secondary graphical analyses was performed with OriginPro software, as described previously (45). Calculated graphs representing the theoretical models were generated by applying the applicable formulae to appropriate input data in OriginPro software.

## RESULTS

### General approach of electrophoresis of fluorophores within supported lipid membranes

TR, NBD and BODIPY were selected as suitable targets for study because, firstly, they are all commonly used fluorescent probes in lipid bilayers and, secondly, they have different chemical and optical properties that could result in a variable susceptibility to quenching or different mechanisms which would make for a good comparison. For example, TR absorbs in the green spectral region (λ ∼ 575 nm) whereas NBD and BODIPY absorb at similar wavelengths in the blue region (λ ∼ 475 nm). Furthermore, TR and NBD are hydrophilic (due to the presence of atoms with different electronegativity, O, N, S) whereas BODIPY is generally considered as hydrophobic (balance between nearby F and N). All three are available to purchase in a form where the fluorophore is covalently tethered onto the headgroup of a lipid. As shown in the chemical structures, the net charge of each fluorophore is zero (**Fig. 2 *A*-*C***) but, when tethered to the negatively-charged lipid DHPE (**Fig. 2 *D***), each molecule has an overall negative charge, *q* = –1*e*, when immersed in an aqueous buffer at neutral pH. Photo-polymerizable lipids were used to generate template patterns consisting of empty 100 × 100 μm squares (**Fig. 2 *F***). Then, a precise concentration of TR, NBD, or BODIPY was incorporated into vesicles comprised primarily from the neutral lipid DOPC (**Fig. 2 *E***), and this solution of vesicles was incubated with the template to form patterned SLBs, termed ‘membrane corrals’ (**Fig. 2 *G***, typically 0.5% fluorescent lipids relative to total lipids, by weight). Fluorophores within these corrals are expected to be mobile and homogenously distributed throughout the membrane (59, 60). When an electric field is applied parallel to an SLB the negatively-charged fluorophores are expected to migrate towards the positive electrode and accumulate at the impenetrable edge of the membrane corrals (**Fig. 2 *H***) (59-61). The balance of forces that work for (e.g., electrophoretic force) and against (e.g., friction and electroosmotic drag) the electric field will determine the fluorophore drift velocity and, eventually, the system will reach a dynamic equilibrium where the fluorophore’s concentration profile across the corral is the result of the competition between E-field-induced drift and random diffusion. To maintain the SLBs under a constant E-field and allow microscopy, samples were maintained within a custom-built flow chamber (see schematic **Fig. 2 *I***). To quantify the self-quenching of each fluorophore, fluorescence lifetime imaging microscopy (FLIM) was used to measure the spatial distribution and corresponding fluorescent lifetimes of fluorophores during electrophoresis experiments. Initial experiments were performed on patterned SLBs containing each fluorophore, to optimize the FLIM image acquisition parameters to ensure the accuracy of fluorescence data collected (accidental photobleaching and singlet-singlet annihilation were minimized, see **Supporting Information** section 1). Fluorescence images and 2-D diffusion measurements confirmed that each fluorophore distributed homogenously throughout the membrane and had high lateral mobility (**Fig. S1**). These preliminary FLIM measurements of patterned SLBs confirmed that high quality data could be acquired for all three fluorophores.

**Figure 2.**
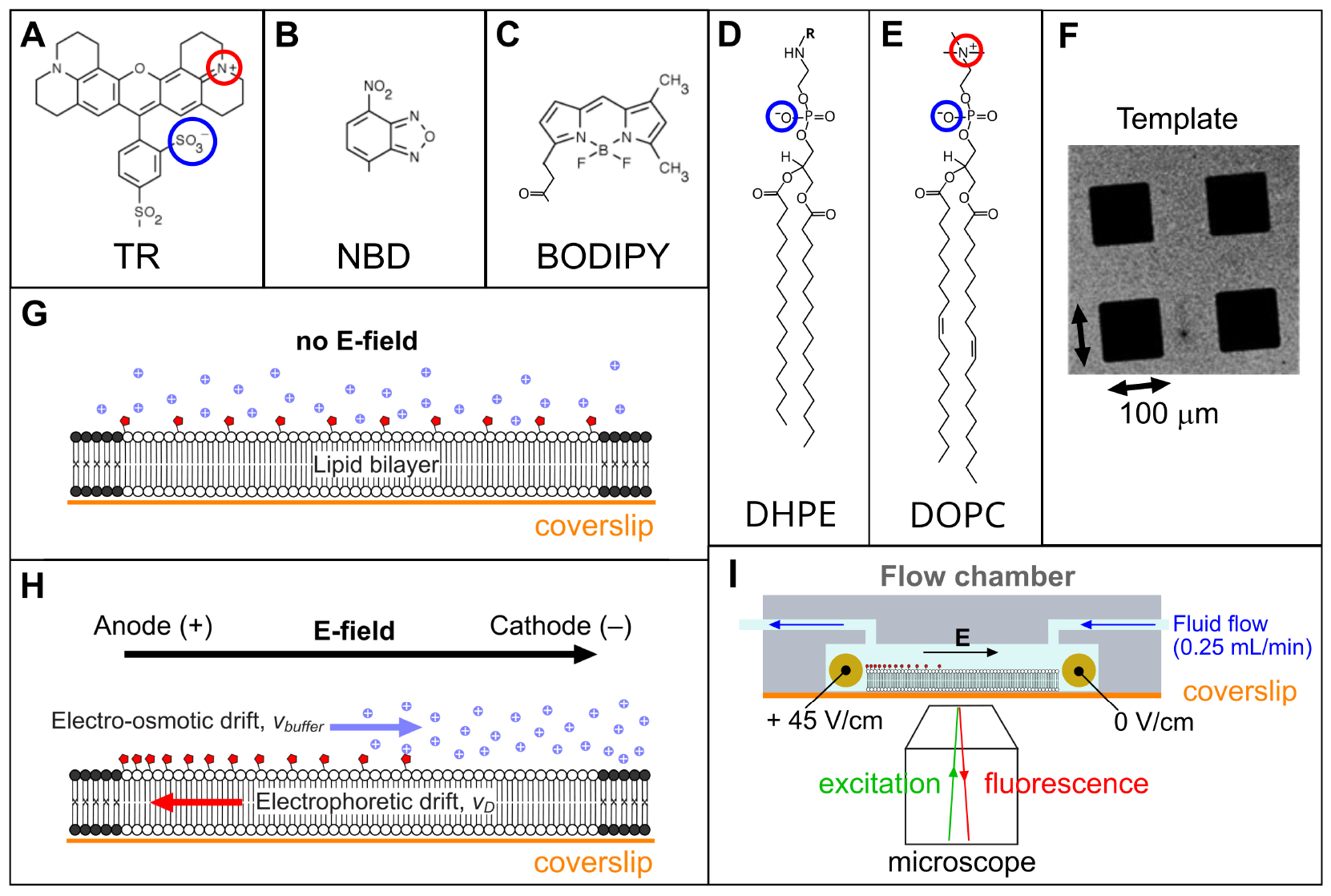
Chemical structures of the fluorophores **(A)** TR, **(B)** NBD, **(C)** BODIPY, and **(D)** the DHPE lipid to which the fluorophores are linked (covalent attachment to replace R). **(E)** The bulk lipid DOPC. **(F)** Example fluorescence microscopy image of the template pattern of Diyne-PC lipids in a micro-array pattern, as generated by photolithography. **(G)** Schematic of a lipid bilayer confined by the barrier of photo-polymerised Diyne-PC (black). In the absence of any electric field, fluorophores (red) will be uniformly distributed in the membrane with a screen of ions (purple) close to the membrane surface. **(H)** As in (G), but with an applied electric field. **(I)** Schematic of the electrophoresis flow cell (not to scale).

### Analysis of FLIM images of fluorophores experiencing quenching

To determine the extent to which different fluorophores are susceptible to concentration quenching, electrophoresis and FLIM was used to analyze membranes containing fluorophores. Corrals of SLBs containing either TR, NBD or BODIPY at a similar initial concentration were compared (0.5% by weight; molar concentrations were considered later) (**Fig. 3**). FLIM images were captured both before electrophoresis and one hour after electrophoresis when a dynamic equilibrium had been established. Before electrophoresis, all three fluorophores had flat homogeneous distributions of fluorescence intensity across the membrane corral (*left panels*, **Fig. 3 *A, D, G***). Qualitatively, the effect of the E-field on the fluorescence intensity was consistent, with all three fluorophores having an asymmetric fluorescence profile with a significantly higher intensity at the left-edge of the corral (*right panels*, **Fig. 3 *A, D, G***). All three fluorophores exhibited over a threefold intensity increase after electrophoresis, to 4.5× the initial intensity for TR, 3.0× for NBD, and 3.8× for BODIPY (**Fig. 3 *B, E, H***). Whilst not a focus of current work, the kinetics of the lipid accumulation could be tracked for each fluorophore and the electrophoretic drift velocity was very similar for all three fluorescent lipids (*v*_*d*_ = 0.31, 0.28, 0.27 μm/s for TR, NBD, BODIPY, see **Fig. S2**) and in agreement with publications on similar systems (50). The estimated concentration of fluorophores accumulated at the positive electrode is calculated later in this work (next section).

**Figure 3.**
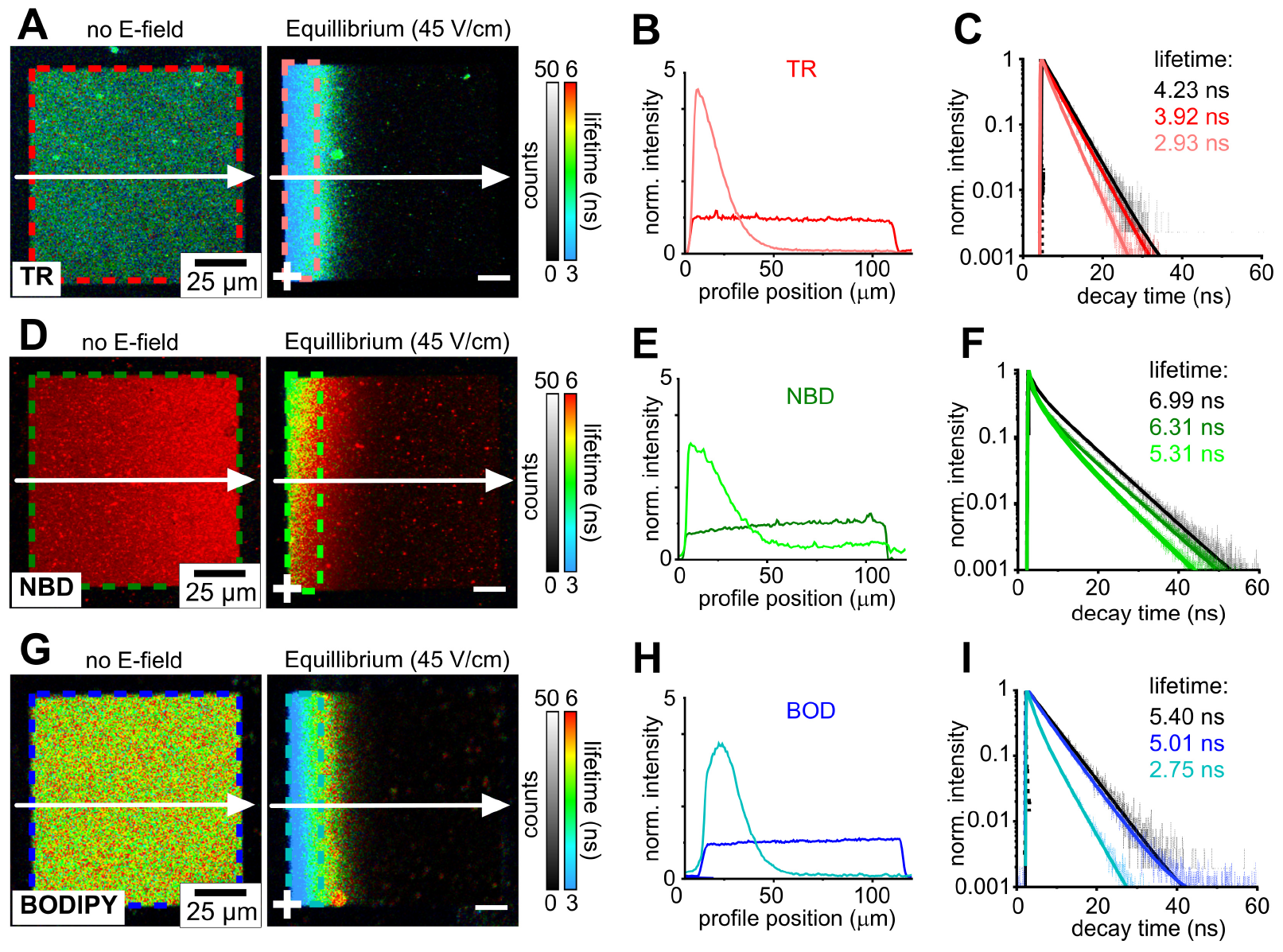
Comparison of the before- and after-electrophoresis states of membrane corrals containing TR, NBD and BODIPY. **(A)** FLIM images of a corral containing 0.5% TR-DHPE (w/w) within a DOPC lipid bilayer, acquired either without any applied E-field (left) versus after the application of a 45 V/cm E-field for 1 hr (right). **(B)** Fluorescence intensity profiles drawn across the membrane (white arrows in (A)), either before (dark red) or after electrophoresis (light red). Both profiles are normalised to 1.0 for the average starting intensity of the corral, in order to compare between samples. **(C)** Fluorescence decay curves representing the membrane corral prior to electrophoresis and after electrophoresis (curves coloured to represent the ROI of the same colour in (A)). For each, the raw fluorescence data (thin lines) was fit to an exponential decay function (bold lines) and the value for mean lifetime extracted from the fit is displayed. The decay curve acquired for a control sample with even lower fluorophore concentration (0.25% TR) is shown for comparison (black curve). **(D)**,**(E)**,**(F)** FLIM images, fluorescence intensity profiles and fluorescence decay curves of a 0.5% (w/w) NBD-DHPE lipid bilayer. **(G)**,**(H)**,**(I)** FLIM images, fluorescence intensity profiles and fluorescence decay curves of a 0.5% (w/w) BODIPY-DHPE lipid bilayer.

For all fluorophores, the fluorescence lifetime was significantly reduced during electrophoresis, as represented by the blue-shift in the false-colour scale for lifetime (*right panels* in **Fig. 3 *A, D, G***). To quantify and compare the quenching between the three fluorophores, lifetimes were determined by fitting the fluorescence decay curves produced from a region-of-interest (ROI) at the left-edge of the corral. It was found that the fluorescence lifetime decreases from 3.92 ns to 2.93 ns for TR, decreases from 6.31 ns to 5.31 ns for NBD, and decreases from 5.01 ns to 2.75 ns for BODIPY (**Fig. 3 *C, F, I***). To quantify the degree of self-quenching for each fluorophore, the quenching efficiency (*QE*) was calculated from the ratiometric reduction in fluorescence lifetime (Eq. 15, see **Methods**). The *QE* observed in the ROI at the left-edge of the corral at electrophoretic equilibrium was calculated as 30.7%, 24.0% and 49.1% for TR, NBD and BODIPY, respectively. Overall, the results in this section show that self-quenching occurs for all three fluorophores and that quenching was induced by increasing their concentration via in-membrane electrophoresis.

### Deeper analysis of quenching efficiency vs. concentration to assess the mechanism

In previous work, we established a method to convert a fluorescence intensity profile into a concentration profile, applying it to TR fluorophores in membrane corrals (45). To do this, the fluorescence intensity is corrected for quenching by quantifying the reduction in fluorescence lifetime (using Eq. 22). The fluorophore concentration can then be obtained using the linear relationship from standard concentration curves of fluorescence intensity (see **Fig. S3-S4**). Here, this method is applied first to TR, then to NBD and BODIPY, so that quenching can be directly correlated with fluorophore concentration, and then compared to theoretical models to understand the mechanism of quenching. Electrostatic repulsions between the negatively-charged fluorophores may limit the final concentrations that can be achieved. Our method simply calculates the fluorophore concentrations that are observed in the system as a result of all effects, irrespective of their origin.

FLIM images of membrane corrals, fluorescence lifetime profiles and concentration profiles for the TR fluorophore are shown in **Fig. 4 *A*-*C***. The calculated molecular concentrations are spatially correlated to fluorescence lifetimes in the FLIM data, so these two types of data can be directly related to each other in a scatter plot to describe the quenching behaviour. **Fig. 4 *D*** shows the resulting *QE* vs *C* curve which combines all data sets obtained for electrophoresis of membrane corrals containing TR. The concentration is displayed as the % mol/mol of fluorophores relative to total lipids (*bottom x-axis*, **Fig. 4 *D***) and also as a number density of fluorophores per 100 nm^2^ to provide a more tangible representation of concentration at sensible length scales (*top x-axis*, **Fig. 4 *D***). From this plot we can make several observations about the quenching behaviour of TR. Firstly, the *QE* vs *C* curves obtained from samples with varying initial concentrations all fall onto one master curve, indicating a highly consistent trend for quenching behaviour. Secondly, the quenching starts to occur immediately after the hypothetical zero concentration, so that even at 0.5% TR (a typical concentration used to probe lipid bilayers) there is a *QE* of 5-10%. Thirdly, the amount of quenching initially increases steeply with concentration before the gradient begins to decrease approaching a *QE* of one, above which it cannot increase by definition. Therefore ∼100% quenching can be expected at higher concentrations above 10% TR.

**Figure 4.**
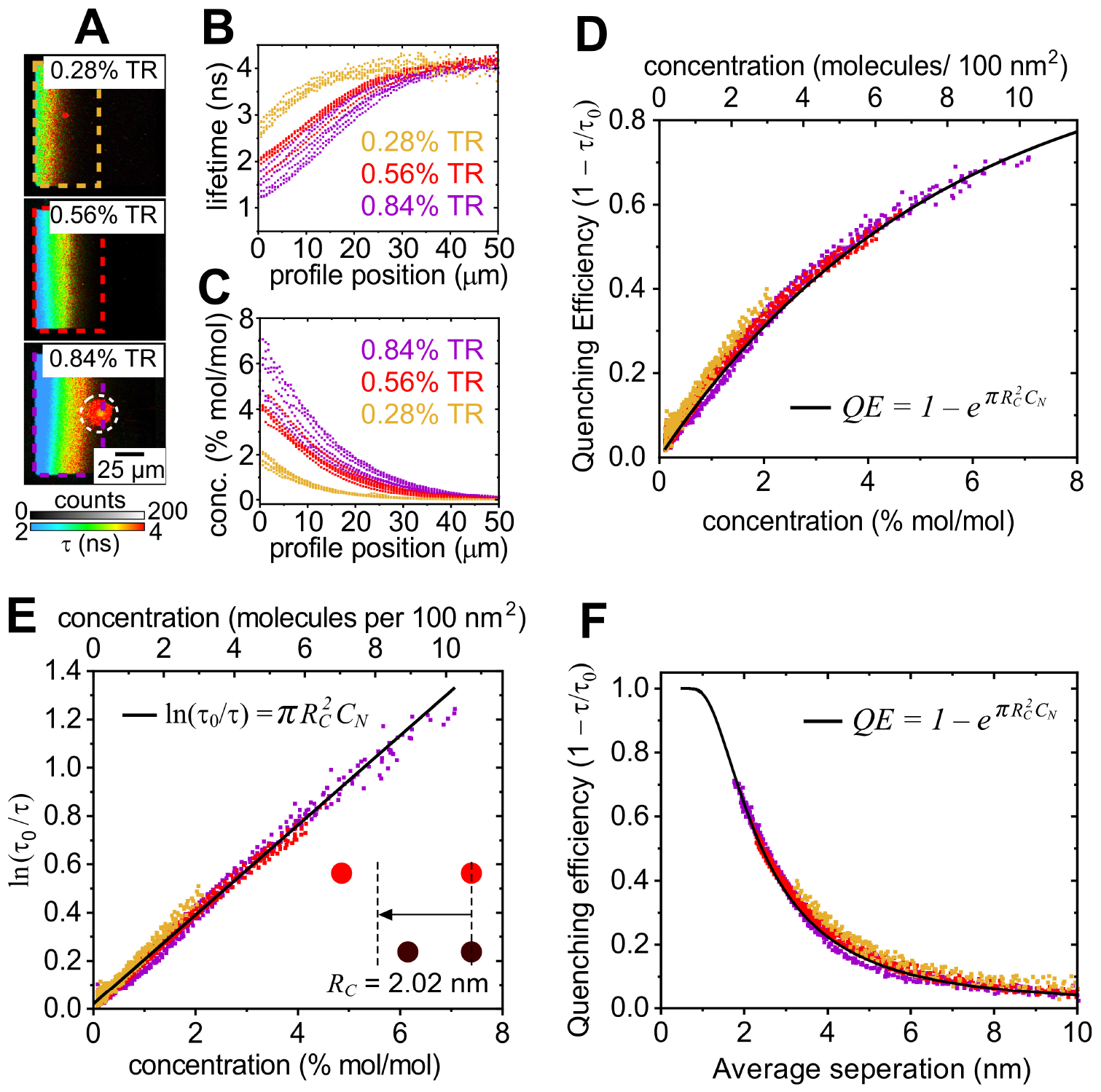
Analysis of quenching efficiency against concentration for a range of TR concentrations. To account for statistical variations between different membrane corrals and improve our precision, the data from multiple corrals for each sample is accumulated (n = 6). **(A)** Example FLIM images of corrals containing an initial concentration of either 0.28, 0.56 or 0.84% (mol/mol) TR at electrophoretic equilibrium. The boxes highlighted by coloured dashed lines are the ROI from which data was analyzed (see **Fig. S4**). The data from these ROI’s were used to generate the corresponding scatter plots in (B)-(F). The circular region indicated by white dashed lines shows an example of a region excluded from the analysis due to a defect in the membrane, likely to be a lipid tubule, as previously reported (45). **(B)** Multiple fluorescence lifetime profiles (overlaid), as obtained from many different corrals. Yellow, red and purple data points represent the profiles obtained from membranes containing 0.28, 0.56 and 0.84% TR, respectively. **(C)** Multiple calculated concentration profiles obtained from the same corrals. **(D)** Quenching Efficiency versus concentration for TR fluorophores in SLBs. The top x-axis was generated from the bottom x-axis (Eq. 1). The solid black line shows the theoretical QE curve calculated using the equation shown (Eq. 18) and the value for R_C_ determined in (E). **(E)** The same data as in (D) plotted as the logarithm of the relative lifetime versus fluorophore concentration. A linear fit (black line) of the data is shown. The gradient 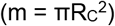of this fit was 12.8 ± 0.1 nm^2^ and R_C_ was calculated as 2.02 ± 0.01 nm. **(F)** The same data as in (D) plotted as QE versus the average separation distance between fluorophores. The solid line shows the theoretical QE curve calculated as in (D).

Next, this data can be compared with the theoretical relationships for transfer-to-trap quenching from the equations derived in the theory section of the **Methods** (Eq. 16-17). A plot of *ln(τ*_*0*_*/τ)* versus *C* was found to have a strongly linear fit (R^2^ = 0.98), the predicted relationship for transfer-to-trap quenching (*black line*, **Fig. 4 *E***). The gradient of this linear fit can be used to calculate the ‘critical radius for trap formation’, *R*_*C*_, a measure of the overall quenching strength of a particular fluorophore that is defined as the separation distance at which two fluorophores have a characteristic (63%) likelihood to associate and form a trap. *R*_*C*_ for TR was calculated as 2.02 ± 0.01 nm, suggesting that two fluorophores must have very low separation distances to form a trap. For example, this is much shorter than the distance at which effective FRET occurs, roughly one-third of the Förster radius for TR (5.73 nm). So far, the quenching behaviour of TR has been considered as a function of the fluorophore concentration but it is also instructive to plot the quenching behaviour as a function of the distance between fluorophores. The average separation distance between fluorophores, *r*, was calculated from the fluorophore number density and *QE* was plotted against this (**Fig. 4 *F***). Following the plot from right-to-left, the *QE* is minimal (<10%) for *r* > 5 nm, before rapidly increasing up to 70% *QE* at an average separation of ∼2 nm (as expected where *r ≈ R*_*C*_). *QE* tends towards 100% at very small *r* where all of the fluorophores become trap sites (at *r* ≪ *R*_*C*_), as observed in the high concentration regime of the *QE* versus *C* plot.

To assess whether the experimental data was consistent with the transfer-to-trap model, theoretical curves for *QE* were generated by using the equations describing this model (Eq. 18) and inputting the desired range of fluorophore concentrations and experimentally determined value of *R*_*C*_ = 2.02 nm. The calculated curves of *QE* versus *C* (*black line*, **Fig. 4 *D***) and *QE* versus *r* (*black line*, **Fig. 4 *F***) were found to be highly consistent with the experimental datapoints for all fluorophore concentrations and equivalent separation distances. Overall, these results show that it is highly likely that TR fluorophores undergo transfer-to-trap quenching, in which fluorophores form non-fluorescent statistical pairs as a probabilistic function of concentration and excitons migrate from excited monomers to these trap sites via FRET.

### Quantitative comparison of quenching behaviour between TR, NBD and BODIPY

A quantitative comparison of the quenching behaviour of all three fluorophores was performed by repeating the analysis described for TR (previous section) for the NBD and BODIPY fluorophores. The *QE* vs *C* curve generated from this analysis is shown in **Fig. 5 *A***. All three fluorophores followed the same overall trend, whereby *QE* increases with *C* and the gradient of the curve becomes increasingly shallow at high concentrations, but the dependence on concentration differs. It is apparent that BODIPY (*blue datapoints*) self-quenches more strongly than TR (*red*) and NBD (*green*), as demonstrated by the steeper gradient of the *QE* vs. *C* curve. For example, 2% (mol/mol) of the fluorophore within a lipid bilayer led to a reduction in the fluorescence lifetime of ∼10% for NBD, ∼30% for TR and ∼50% for BODIPY. This implies that BODIPY fluorophores form trap sites more readily than TR or NBD. Next, to assess whether or not all three fluorophores exhibit transfer-to-trap quenching, each data set was plotted as *ln(τ*_*0*_*/τ)* versus *C* and fitted to the predicted linear relationship. For all three fluorophores, the experimental data was strongly correlated to the theoretical model with a high quality fit over all concentrations (**Fig. 5 *B***), suggesting that they undergo the same quenching mechanism. To compare the relative strengths of quenching, the gradients of these linear fits were determined and *R*_*C*_ values were calculated as 2.70 nm, 2.02 nm and 1.14 nm for BODIPY, TR and NBD, respectively. Therefore, we find that the quenching strength of BODIPY is 1.3 times that of TR, which is 1.8 times that of NBD.

**Figure 5.**
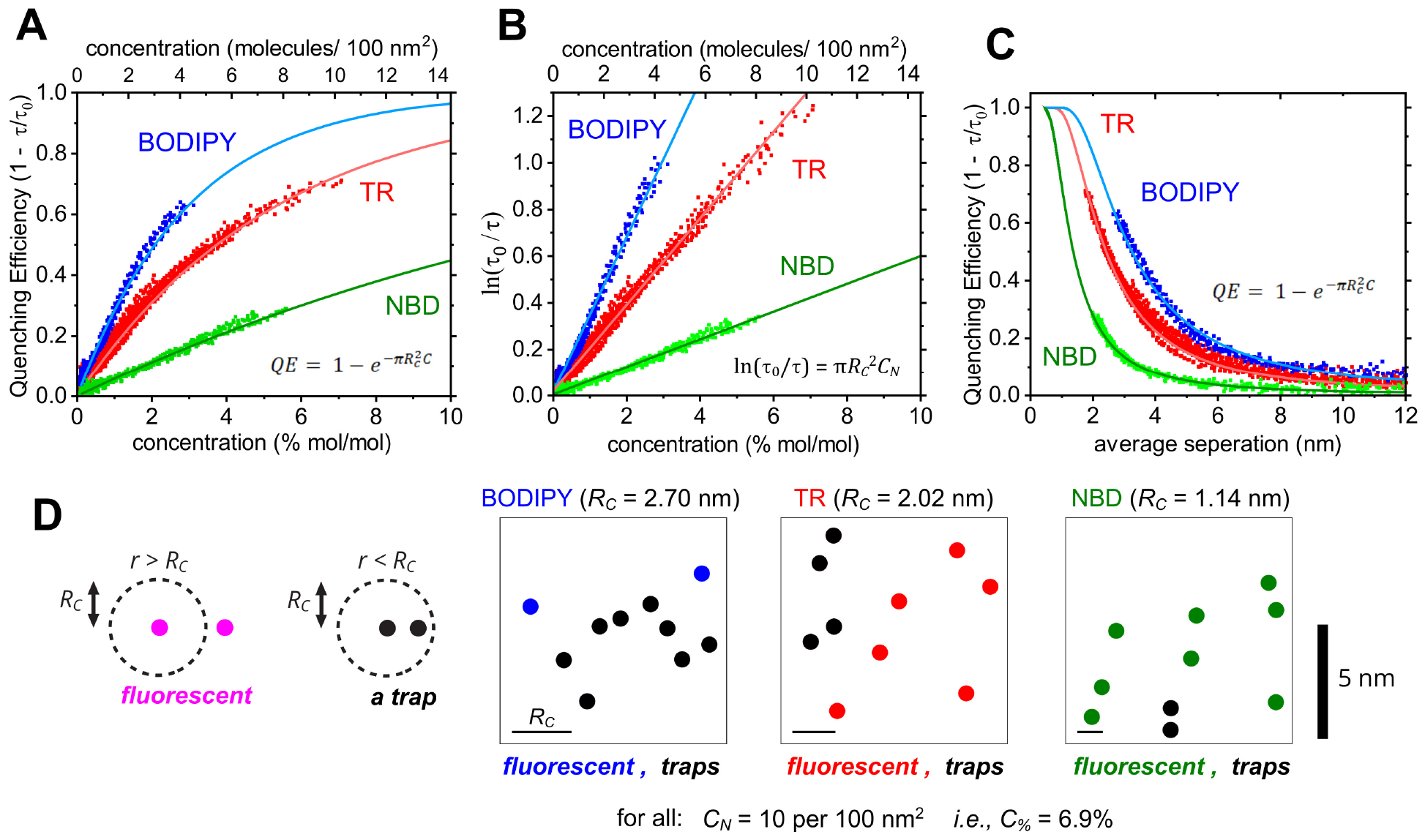
Graphs comparing the quenching relationships between TR, NBD and BODIPY. **(A)** Quenching Efficiency versus concentration for each fluorophore type. The solid lines show the theoretical QE curves calculated using the equation shown (Eq. 18) and the values for R_C_ determined in (B). **(B)** The same data as in (A), plotted as the logarithm of the relative lifetime versus fluorophore concentration. Linear fits (solid lines) of the equation shown were used to obtain the critical radius for trap formation, R_C_. The gradient 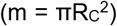 of each fit was 23.0 ± 0.1, 12.8 ± 0.1 and 4.11 ± 0.02 nm^-2^ for BODIPY, TR and NBD, respectively, leading to R_C_ = 2.70 ± 0.01, 2.02 ± 0.01 and 1.14 ± 0.01 nm. **(C)** The same data as in (A) plotted as QE versus average separation distance between fluorophores. The solid lines show the theoretical QE curves calculated as in (A). **(D)** Representations of a 10 × 10 nm region of lipid bilayer containing either TR, NBD or BODIPY, showing fluorophores that are either fluorescent (in colour) or acting as traps (black), according to the R_C_ values calculated in (B). This is an oversimplification because R_C_ is not a threshold value but instead a relative measure of the propensity to quench. Each box contains the same concentration of fluorophores but different numbers of traps exist due to the different R_C_.

To consider how these trends relate to distances between molecules, *QE* vs *r* plots were generated for each fluorophore (**Fig. 5 *C***). As expected, following the trend lines from right-to-left, BODIPY begins to quench at higher separations than TR and NBD, and BODIPY reaches a *QE* of 50% at ≈3.3 nm compared to ≈2.4 nm for TR and ≈1.4 nm for NBD. At decreasing separation distances, towards zero, the *QE* continues to increase and may be expected to saturate at *QE* = 1 for very low separations. Theoretical quenching curves generated using the fitted values for *R*_*C*_ show strong correlation to the *QE* versus *C* data (*solid lines*, **Fig. 5 *A*** and ***C***). Overall, the strong correlation between experimental data and the theoretical relationship representing quenching by statistical pairs (traps) and excitation energy transfer throughout the membrane (transfer-to-trap) is good evidence that this mechanism does indeed occur for all three fluorophores. A final consideration of the accuracy of the trends for fluorescence quenching can be made now that *R*_*C*_ values have been determined for each fluorophore. A simplified mathematical model of quenching has been used so far that assumes FRET always occurs, so that the equations could be solved and *R*_*C*_ quantified (see theory section of the **Methods**). The simplified model for quenching (*P*_*FRET*_ = 1) was compared with the exact model (*P*_*FRET*_ varies) by calculating the fluorescence intensity and lifetime as a function of concentration using the appropriate equations and *R*_*C*_ values (see **Fig. S5**). There was only an observable discrepancy at low fluorophore concentrations (when *P*_*FRET*_ drops) and this caused only minor differences in fluorescence intensity and lifetime. This is logical because at low concentrations there is little quenching anyway and at high concentrations the simplified model is very close to the exact model. This confirms that quenching is dominated by the fraction of traps, rather than the FRET efficiency, for all three fluorophores.

To visualize the number of traps sites expected to exist based on the calculated critical radii, a schematic was drawn to show the same concentration of fluorophores for each fluorophore in a random distribution with trap sites indicated based on the calculated *R*_*C*_ values (**Fig. 5 *D***). At the ∼7% concentration shown, it is evident that the majority of BODIPY molecules would exist as traps, as compared to very few traps for NBD and an intermediate number of traps for TR.

## DISCUSSION

### Mechanism of quenching: transfer-to-trap and statistical pairs

From 1950-2000 there were several experimental and theoretical investigations of randomly-distributed pigments as a model system for photosynthesis, particularly before the biological light-harvesting structures were known to be protein-pigment complexes. These studies proposed that the concentration-induced fluorescence quenching of chlorophyll pigments could arise either due to the occurrence of long-lived physical aggregates at very high pigment concentrations (over 100 mM) or due to ‘statistical pairs’ via a transfer-to-trap mechanism at lower pigment concentrations (at 2-100 mM) (5-7, 31, 32, 34, 36, 42, 43, 62) (see **Fig. 1** and **Methods** theory section). A small number of recent studies have provided evidence for quenching by statistical pairs of chlorophylls using Monte Carlo simulations (37), molecular dynamics simulations (39) and mathematical modelling (35). Over the last decade, there has been great focus on understanding the potential mechanisms of quenching that occur in LH proteins, because quenching is thought to be crucial as part of a protective process for dissipating energy under high-intensity sunlight (15). However, there is little agreement on the identity of the quencher in LH proteins, possibly due to the high complexity of the system (16-25). Therefore, we decided that our newly-optimized experimental technique to generate concentration gradients of fluorophores and analyse them with an advanced form of microscopy would be an ideal platform to test for agreement or disagreement with previously proposed theoretical mechanisms, and, furthermore, to compare multiple fluorophores that have different chemical structures. In this paper, we quantified the fluorescence intensity and kinetics of three organic fluorophores: TR, NBD and BODIPY. The mechanism of self-quenching for all three fluorophores was found to be a combination of the concentration-dependent formation of ‘trap sites’ (statistical pairs) and the transfer of excitation energy between fluorophores until they reach these sites, i.e., ‘transfer-to-trap’. The evidence for this and its wider implications are highlighted below.

Fluorophores undergoing electrophoresis within lipid bilayers were observed using fluorescence microscopy, and the fluorescence lifetimes were analysed, allowing the local concentration of fluorophores to be cross-correlated to their quenching (**Fig. 3-5**). The extent of quenching for TR, NBD and BODIPY was found to increase with fluorophore concentration before tending towards unity at very high concentrations (**Fig. 5**). For all three fluorophores, the mathematical expression representing the transfer-to-trap theory was in excellent agreement with the experimental data suggesting that this is the correct explanation for quenching. The important implication here is that self-quenching depends upon ‘statistical pairs’ of fluorophores which do not require any interaction between their ground states (7, 9, 31, 32). In other words, traps may occur simply due to the temporary proximity of one fluorophore to another at a given concentration and do not require chemical/ physical interactions or a specific molecular configuration. This may explain why such quenching behaviour appears to be a general phenomenon of both natural and synthetic pigments that do not share chemical similarities and thus have no reason to self-associate in similar manners.

### Comparison of the quenching strength of three different fluorophores

Whilst the self-quenching of all three fluorophores could be explained by a transfer-to-trap model, there were significant differences in the extent of this quenching (**Fig. 3, Fig. 5**). The critical radii for trap formation, representing the relative quenching strengths, were found to be *R*_*C*_ = 2.70, 2.02 and 1.14 nm, for BODIPY, TR and NBD, respectively. These *R*_*C*_ are in close agreement to those found in previous investigations of self-quenching of NBD (9) and BODIPY (10) and give us confidence in the accuracy of our analyses. To our knowledge, the *R*_*C*_ of TR has not been previously reported. To summarize: BODIPY undergoes concentration-dependent quenching most strongly, followed by TR at an intermediate level, and then NBD at a lower level. Previous studies reported that the *R*_*C*_ of chlorophyll is ≈ 1 nm (32, 37) and therefore a relatively weak quencher similar to NBD. Despite our characterisation of the quenching strengths of TR/ NBD/ BODIPY, it is not explicitly clear why different fluorophores have different critical radii. We note that the potential for trap formation does not follow the same trend as the potential for FRET represented by their Forster radii, *R*_*0*_ (5.41, 5.73, 3.32 nm for BODIPY, TR, NBD). NBD has both the lowest *R*_*C*_ and *R*_*0*_ so we can speculate that the relatively poor energetic coupling between an excited-state NBD and a ground-state NBD may hinder both FRET and trap formation. In contrast, BODIPY has a lower *R*_*0*_ but a higher *R*_*C*_ than TR, therefore there must be something beyond the Förster coupling parameters that affects trap formation.

One possibility is that chemical differences may cause some fluorophores to have a greater potential for attractive interactions than other fluorophores, and may form long-lived (physical) dimers or larger aggregates. As TR, NBD and BODIPY are negatively-charged (in water at pH 7) they will experience electrostatic repulsion, but this must be balanced against any attractive interactions such as those between the nonpolar tailgroups of the lipids that the fluorophores are tethered to. BODIPY has been reported to aggregate at high concentrations as a result of hydrophobic interactions between planar moieties of the molecule (41, 63). Similar face-to-face ‘H-type’ dimers (or H-aggregates) have also been suggested to occur for chlorophylls at very high concentrations (37, 42). Long-lived dimers could explain the greater quenching strength (greater *R*_*C*_) of BODIPY relative to TR/ NBD, i.e., there could be some combination of the transfer-to-trap model and the aggregate model for BODIPY. Absorbance spectroscopy could identify (or rule out) the presence of long-lived physical dimers/ aggregates (42, 64), however, absorption spectroscopy of SLBs on glass coverslips was not possible with our instrumentation. Whilst our experiments do rule out the presence of ground-state dimers, we highlight that the transfer-to-trap theory with statistical pairs was sufficient to model the data.

### The molecular identity of the trap site and the energy dissipation pathway

The transfer-to-trap model defines the relationship between fluorophore concentration and their tendency to form trap sites, however, it does not specify the photophysical mechanism for quenching, i.e., how energy is dissipated once an excited state reaches a trap site. Two photophysical mechanisms of energy dissipation have been proposed in previous studies of quenching of fluorophore pairs. The first mechanism is quenching by a ‘dark state’, where the trap site is a pair of fluorophores that undergo ‘excitonic splitting’ to form one super-radiant state and one dark state (37, 65, 66). Fluorescence is disallowed from the dark state, therefore, non-radiative decay will occur to dissipate the energy as heat. Such split excitonic states have been suggested to occur for H-aggregates and excimers of fluorophores (37, 41, 63, 67). Alternatively, a second mechanism, is quenching via a charge-transfer state, sometimes called photoinduced electron transfer. In this mechanism, the trap site is a pair of fluorophores that form a charge-transfer (ion-pair) state after receiving excitation energy (39, 40, 62). This charge transfer state undergoes rapid recombination to regenerate the ground state and in the process dissipate the energy as heat (43, 44). Both of these proposed mechanisms have a unique spectral signature, however, this is technically challenging to observe experimentally because of the transient nature of the molecular species. Whilst the molecular identity of the quencher was not investigated in the current study, we would like to highlight that it would be an interesting topic for future studies. Transient absorption spectroscopy could resolve changes in the nature of the excited state at the very short timescales involved (21, 68, 69) and Stark fluorescence spectroscopy can identify charge-transfer states (69, 70). Completing our understanding of both the occurrence of the traps and their molecular identity will result in a more holistic understanding of how concentration-induced fluorescence quenching can occur in model systems and would have direct relevance as potential mechanisms of quenching in biological LH pigment-protein complexes. Future studies could use the electrophoresis/ FLIM approach as a novel method to generate high concentrations of LH proteins and quantify the resultant quenching.

## SUPPORTING INFORMATION

Supporting Information is available - including supplementary text, five additional figures (**Fig. S1-S5**) and three tables of data (**Table S1-S3**).

## AUTHOR CONTRIBUTIONS

P.G.A. and S.A.M conceptualised the project. S.A.M. developed theory, prepared samples and performed FLIM analyses. Y.K. prepared the template patterns for microscopy. P.G.A., S.D.E, S.D.C. and K.M. provided supervision over the course of the project and acquired funding. S.A.M. and P.G.A. wrote the first draft of the manuscript. All authors contributed to revising the manuscript.

## ACKNOWLEDGEMENTS

S.A.M. was supported by a PhD studentship from the Biotechnology and Biological Sciences Research Council UK (BBSRC) (BB/M011151/1). Y.K. and K.M. were supported by the Japan-UK Research Cooperative Program (JPJSBP120195707) and a Grant-in-Aid for Scientific research (Kakenhi) (No. 19H04725 and 21KK0088) from Japan Society for the Promotion of Science (JSPS). S.D.E. was supported by the National Institute for Health Research infrastructure at Leeds and by the EPSRC (EP/W033151/1). S.D.C. was supported by grants from the Engineering and Physical Sciences Research Council UK (EPSRC) (EP/R043337/1, EP/R03608X/1 and EP/J017566/1). P.G.A. was supported by a University Academic Fellowship from the University of Leeds and a grant from the EPSRC (EP/T013958/1). The PicoQuant FLIM instrument at Leeds was acquired with funding from the BBSRC (BB/R000174/1).

