## Supplementary Information for "Evidence for a transfer-to-trap mechanism of fluorophore concentration quenching in lipid bilayers"

for the article:

### SI 1. Confirming the membrane quality: structural contiguity and lateral lipid diffusion

The FLIM instrumentation was set up as described in the **Experimental Methods** section (main text), changing the excitation laser and emission filters used so that they were appropriate for the fluorophore. For TR excitation at 561 nm and collection between 590-650 nm was used, whereas, for NBD and BODIPY excitation at 485 nm and collection 500-540 nm was used. FLIM images of lipid membranes containing 0.5% (w/w) NBD, TR, and BODIPY are shown in **Figure S1A**. In each case, fluorescence is restricted to well-defined square patterned membranes, with minimal signal located on the surrounding template. The fluorescence within each membrane is largely homogeneous across the corral (40-50 counts/pix for all fluorophores), with a few visible bright spots that may represent non-ruptured vesicles that are loosely adsorbed onto the membrane. Overall, these membranes were highly reproducible, with minimal variation in intensity and quality across multiple preparations.

To confirm that the liposomes had ruptured to form well-connected membranes containing mobile fluorophores (and not merely adsorbed into the template without rupturing), fluorescence recovery after photobleaching (FRAP) experiments were performed to monitor the diffusion of lipids over time. For each sample, a circular area (with a bleached radius, $R_{bleach}$, ranging from 20-30 μm) of membrane was deliberately photobleached using intense white light. Immediately after photobleaching, a FLIM timelapse of images (**Figure S1B**) was obtained to monitor the diffusion of "non-bleached" fluorophores into the bleached area. The intensity of the fluorescence recovery in the bleached spot was monitored for each timepoint (each time point is the accumulation of photons in a 16 s period) to plot a fluorescence recovery curve (**Figure S1C**). A mono-exponential fit, $F = F_0(1 - e^{-kt})$, was used to obtain the rate constant $k$ and then the "doubling time", $\tau = ln(2)/k$, for each sample. From this, the diffusion constant, $D = 0.22 \times R_{bleach}^2/\tau$, was calculated. To calculate the mobile fraction, images of the corral before photobleaching and after photobleaching were analysed. Briefly, the intensity of the "bleached" region was compared to the intensity of a "non-bleached" region throughout the FRAP experiment. For a mobile fraction of 100% the two regions would have an equal intensity after the system has been allowed to reach an equilibrium. A summary of FRAP experiments on patterned bilayers is shown in **Table S1.** For each fluorophore, the diffusion of the lipids was compared in both patterned bilayers (lipid bilayers ruptured into a 100 μm square template) and in "infinite" bilayers (lipid bilayers ruptured onto non-patterned glass) to ensure that the Diyne-PC template did not adversely affect the lipid mobility. The calculated diffusion constant was similar for all fluorophores in both patterned and non-patterned bilayers (ranging from 1.75 to 2.27 $\mu m^2/s$) and the mobile fraction was consistently high across all samples (>94% for all fluorophores). Overall our results are consistent with the values for the diffusion constant and mobile fractions of similar lipid-tagged fluorophores reported in other studies[1, 2], and a strong indication that any interactions that the lipid-tagged fluorophores may have with either the glass substrate or the Diyne-PC template do not hinder the lateral diffusion of lipids or the electrophoretic effect. In addition, the mobility, $\mu_{FRAP} = D/k_BT$, was calculated for each fluorophore (ranging between 4.2 to 5.5 $ms^{-1}N^{-1}$). The mobility is a measure of how a particle responds to an applied force, and will be used to quantify the electrophoretic effect investigated, see **SI 2**.

The accidental photobleaching of fluorophores by the excitation laser could have the effect of altering the concentration of "active fluorophores", which would be undesirable and lead to misinterpretation of fluorescence microscopy data. Photobleaching will increase with laser power and acquisition time. Therefore, the photobleaching of the fluorescent probes was quantified and minimized, as described in reference [3]. Similarly, singlet-singlet excited state annihilation effects can distort fluorescence measurements by reducing the values measured for fluorescence lifetime. The probability of annihilation effects increases with laser power or, more accurately, the energy per pulse per unit area (illumination area on the sample surface). Therefore, a laser power dependence measurement series was made to quantify annihilation effects for these fluorescent probes, as shown in reference [3]. As found in that publication, the optimal laser fluence for this instrument and these type of samples was found to be ~0.012 mJ $cm^{-2}$ with a laser spot diameter of ~800 nm (10 MHz repetition rate, pulse widths of ~70-100 ps for the 485 nm laser and 561 nm laser). With these laser settings, there were no detectable annihilation effects and photobleaching over the course of a standard FLIM image acquisition of 25 frames (80 s of exposure) was below 2%, for all fluorescent probes used.

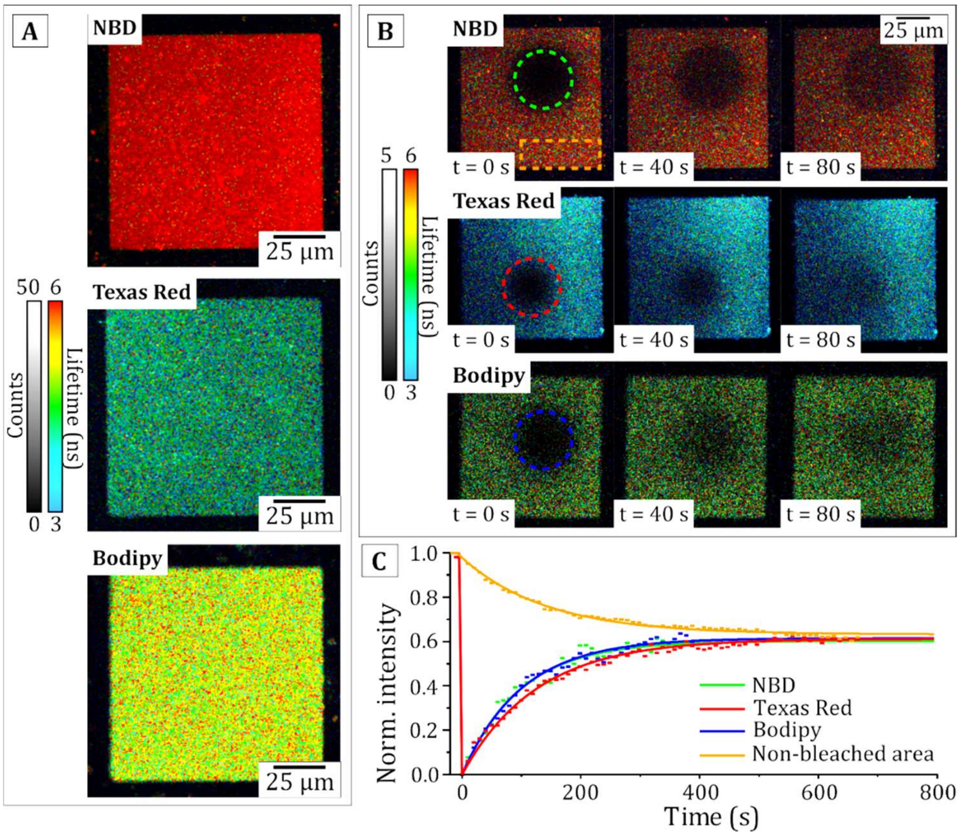


***Figure S1:*** *FLIM images of TR-, NBD- and BODIPY-containing lipid bilayers and Fluorescence Recovery After Photobleaching (FRAP) experiments confirming that the fluorophores are mobile and a suitable target for electrophoresis.* ***(A)*** *Example images of DOPC lipid bilayers containing 0.5% (mol/mol) of either NBD (top), TR (middle) or BODIPY (bottom) formed as 100 µm wide corrals within the DiynePC polymer templates. These are all set to the same fluorescence lifetime scale of 3-6 ns to allow a visual comparison of the relative differences in lifetime.* ***(B)*** *FRAP experiments for the patterned lipid bilayers, showing the mobility of each type of fluorophore.* ***(C)*** *FRAP recovery curves showing the fluorescence intensity in the dashed bleached regions in (b), normalised to the intensity of the bilayer prior to photobleaching (colours as labelled on legend). The orange curve shows the fluorescence intensity decrease in the orange, box region in (b), showing that the intensity in the patterned membrane decreases due to there being a finite number of non-bleached fluorophores available to diffuse into the bleached region. The FRAP data on TR is reproduced from Meredith et al. 2023 [3] (the other data is new).*

| | TR | | NBD | | BODIPY | |
|---|---|---|---|---|---|---|
| | Patterned bilayer | Infinite bilayer | Patterned bilayer | Infinite bilayer | Patterned bilayer | Infinite bilayer |
| D ($\mu m^2 s^{-1}$) | 2.27 ± 0.59 | 1.95 ± 0.20 | 2.13 ± 0.18 | 1.99 ± 0.20 | 2.09 ± 0.39 | 1.75 ± 0.17 |
| $\mu_{FRAP}$ ($ms^{-1}N^{-1}$) | 5.49 ± 0.14 | 4.71 ± 0.48 | 5.15 ± 0.43 | 4.81 ± 0.48 | 5.05 ± 0.94 | 4.23 ± 0.41 |
| Mobile % | 95.5 ± 2.5 | 96.6 ± 3.6 | 94.7 ± 2.3 | 97.0 ± 5.2 | 94.0 ± 1.8 | 95.3 ± 7.2 |

***Table S1:*** *Summary of FRAP experiments comparing patterned lipid bilayers (SLBs in a 100 µm square corrals) versus non-patterened lipid bilayers (SLBs on piranha-cleaned glass coverslips). Overall, the similarity of the results between different substrates indicates that the prescence of the polymerized lipid (Diyne-PC) template has no significant effect on the mobility of lipids close to the centre of the membrane.*

### SI 2. Full calculation of electrophoretic drift velocity for all TR, NBD and BODIPY

A series of time-lapse measurements are shown for corrals of SLBs containing TR/DOPC (**Figure S2A-C**). The drift velocity for TR was found by fitting a straight line (*blue line,* **Figure S2D**) to the moving edge displacement plot and found to be 0.28 ± 0.01 μm/s (fitted value ± uncertainty). The analysis of TR was recently published in another paper from our group [3]. The analysis process was repeated for NBD and BODIPY and the fluorophore mobility was calculated (**Figure S2E**). A summary of the calculated kinetic parameters for all three fluorophores is shown in **Table S2**. To determine the contribution of electroosmotic drag to the mobility of fluorophores, the electrophoretic mobility, $\mu_{EP} = V_d / qeE$, was calculated and compared to the FRAP mobility, $\mu_{FRAP}$, for each NBD, TR and BODIPY (from **Figure S1** and **Table S1**). The ratio of the two mobilities, $\alpha = \mu_{EP}/\mu_{FRAP}$, was calculated for each fluorophore to describe the reduction in the mobility due to electroosmotic drag. The reduction factor was found to be 0.83 ± 0.10, 0.70 ± 0.04 and 0.74 ± 0.02 for NBD, TR and BODIPY respectively. These values are similar to the values for α found in other studies of electrophoresis of lipid tagged fluorophores [2], and show that all three fluorophores are affected by electroosmotic drag to some degree. The different reduction factor for each fluorophore is likely due to the amount that each fluorophore protrudes into the aqueous buffer, for example, a large, bulky molecule will experience more friction and drag from the ions close to the membrane. Characterising the kinetics of electrophoresis in this manner shows that it is possible to control the re-organisation of fluorophores, with similar success for each fluorophore.

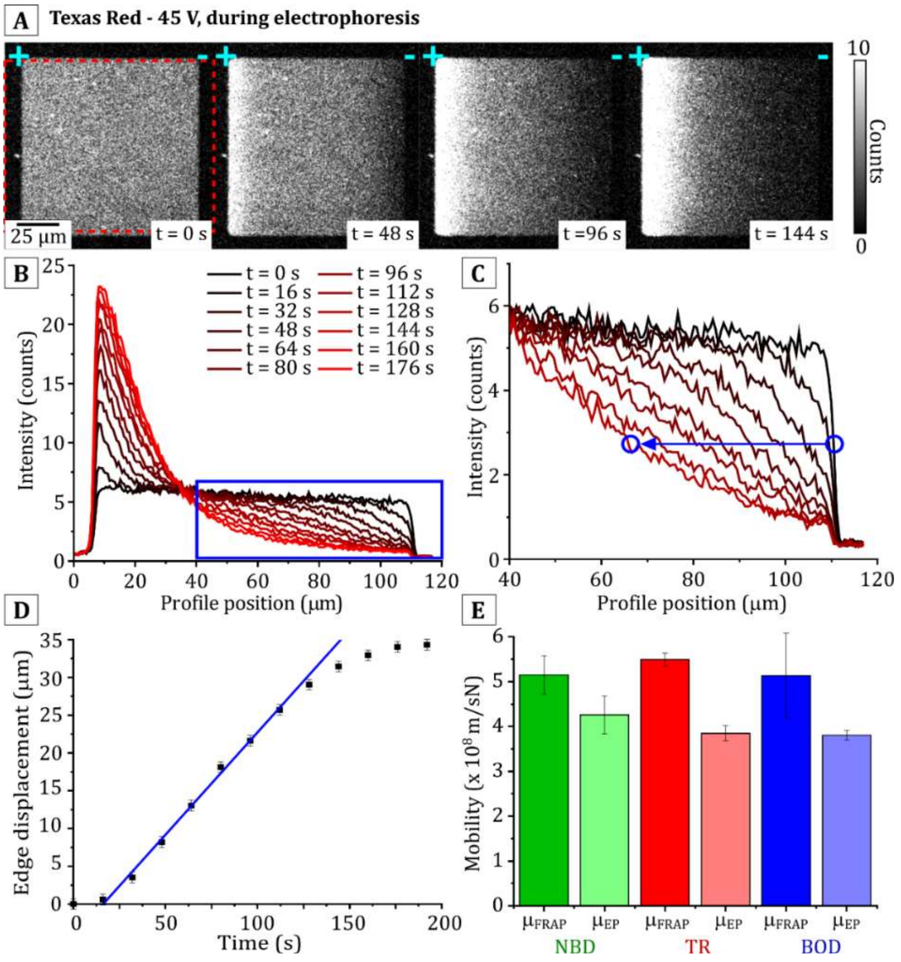


***Figure S2:*** *Timelapse analysis of the drift velocity of TR, NBD and BODIPY.* ***(A)*** *Timelapse series of FLIM images (intensity-only) of a bilayer containing 0.5% (w/w) TR-DHPE after the application of an electric field (45 V/cm).* ***(B)*** *Average intensity profiles measured in the red, dashed box region in panel (a). Black-to-red lines represent increasing timepoints separated by 16 s intervals.*

***Figure S2 (caption continued): (C)*** *A higher magnification portion of panel (b), focusing specifically on the "moving edge" at the right side of the patterned bilayer. The edge midpoint (position of the half-maximum intensity) is measured (blue circle) for each timepoint.* ***(D)*** *Graph showing the displacement of the moving edge of fluorescence with increasing timepoints. The drift velocity, $V_{drift}$, is obtained from a linear fit (blue line).* ***(E)*** *Graph showing the fluorophore mobility as measured by FRAP, $\mu_{FRAP}$, and as measured through by electrophoresis, $\mu_{EP}$ for NBD (green), TR (red), and BODIPY (blue). Panels A-D are reproduced from Meredith et al. 2023 [3].*

| Fluorophore | $\mu_{FRAP}$ (×10$^8$ m/sN) | $v_d$ (μm/s) | q (e) | E (V/μm) | $\mu_{EP}$ (×10$^8$ m/sN) | α |
|---|---|---|---|---|---|---|
| *TR* | 5.49 ± 0.14 | 0.31 ± 0.03 | -1 | 0.0045 | 3.85 ± 0.17 | 0.70 ± 0.04 |
| *NBD* | 5.15 ± 0.43 | 0.28 ± 0.01 | -1 | 0.0045 | 4.26 ± 0.42 | 0.83 ± 0.10 |
| *BODIPY* | 5.14 ± 0.94 | 0.27 ± 0.08 | -1 | 0.0045 | 3.81 ± 0.11 | 0.74 ± 0.02 |

***Table S2:*** *Summary of the calculation of kinetic parameters for diffusion of TR, NBD and BODIPY.*
*$\mu_{FRAP}$ is the Einstein mobility of each fluorophore as in Table S1;*
*$v_d$ is the drift velocity of each fluorophore as estimated in Figure S2D;*
*q is the charge per molecule at neutral pH in units of e (elementary charge, $1.602\times10^{-19}$ C);*
*E is the electric field strength applied (measured during the experiment with a voltmeter);*
*$\mu_{EP}$ is the electrophoretic mobility of each fluorophore.*
*α is the ratio of electrophoretic and Einstein mobilities, $\alpha = \mu_{EP}/\mu_{FRAP}$.*
*For fluorophores measured in a previous study of TR electrophoresis [2], the electrophoretic mobility was found to be 40% lower than the Einstein mobility due to the effect of electroosmotic drag (i.e., α ≈ 0.6). Here we found that $\mu_{EP}$ was 20-30% lower (i.e., α ≈ 0.7-0.8).*

**SI 3. Conversion of raw fluorescence intensity data to molecular concentration by using a standard curve and correcting for the effects of quenching**

It is important for our analysis of quenching to be able to quantify the concentration of the molecules of interest. In a recent study [3], we showed that it is possible to convert experimentally-measured values for fluorescence intensity into the molecular concentration of the fluorescent molecule (Texas Red). Similar analysis was performed on NBD and BODIPY, in addition to TR, as described below.

Supported lipid bilayers (SLBs) were formed on hydrophilic glass, containing different fluorophore concentrations in a range from 0.25-1.5% weight of fluorophore relative to weight of total lipids, of either TR, NBD or BODIPY. We later realized that units of "mole-to-mole" for concentrations were required for the theoretical modelling, therefore, each weight-to-weight concentration was converted to a mole-to-mole percentage before performing the analysis. These SLBs (without any template pattern) acted as simple control samples, due to their high reproducibility, high throughput, and the ability to accurately prepare a defined concentration of fluorophores. FLIM analysis was performed on the SLBs using the same image acquisition parameters as described for electrophoresis measurements on membrane corrals so that the fluorescence intensities and lifetimes were perfectly comparable. FLIM images for these samples are displayed in **Figure S3A-C** and show that the fluorescent lipids were homogeneously distributed across the SLB, as expected. The average fluorescence intensity and average fitted lifetime was calculated for each image. The raw data for the observed fluorescence intensity (datapoints labelled *F* in **Figure S3D-F**) was found to increase non-linearly with fluorophore concentration for TR, NBD and BODIPY, due to the increasing effect of quenching that occurred at higher concentrations. Concurrent with this, the fluorescence lifetime was found to gradually decrease as fluorophore concentration increased (datapoints labelled $\tau$ in **Figure S3D-F**)

In the absence of quenching, the fluorescence intensity would increase linearly with concentration because, logically, with 2× the density of molecules there will be 2× the number of excited states generated for any given excitation laser pulse and, thus, 2× the number of photons of fluorescence expected. This would be a simple relationship from which to estimate the fluorophore concentration, however, this is not observed in the raw data because the fluorescence intensity is somewhat reduced as a result of quenching. As defined in the main text *Eq. 15*, Quenching Efficiency, $QE = 1 - \tau/\tau_0$, so the reduction in fluorescence lifetime is a measure of quenching. The fluorescence lifetime ($\tau$) measured in a FLIM image was used to calculate the relative change in lifetime, $\tau/\tau_0$, where $\tau_0$ is the lifetime in the absence of any quenching (here, $\tau_0$ was defined as the lifetime measured for the fluorophore within an SLB at 0.25%). This ratio $\tau/\tau_0$ was then used to calculate the "non-quenched" fluorescence intensity ($F_0$) from the raw fluorescence intensity ($F$), according to the expression derived in the main text:

$$F_0 = F \cdot e^{\left[2\ln\left(\frac{\tau_0}{\tau}\right)\right]} \qquad \text{Main text Eq. 22}$$

The result of this calculation is shown in **Table S3** and plotted for each fluorophore in **Figure S3D-F** (datapoints labelled *F0*). As expected from our logic that fluorescence must be proportional to concentration in an ideal system, the "corrected fluorescence intensity" $F_0$ does indeed appear to have a linear increase with the concentration and was successfully fit to a straight line, $F_0 = mC + Y_0$, where $m$ is the fitted gradient and $Y_0$ is the y-intercept ($R^2 > 0.99$ for all three datasets). The finding that there is a good linear fit is excellent evidence that the method to estimate the non-quenched fluorescence is accurate. The relationships between the corrected fluorescence intensity and fluorophore concentration were found to be $F_0 = 168.3 \times C_{TR} + 4.8$, $F_0 = 73.0 \times C_{NBD} + 9.7$, and $F_0 = 168.8 \times C_{BODIPY} + 15.3$ for TR, NBD and BODIPY respectively. The fact that $Y_0$ is not equal to zero for all three fluorophores is likely to be due to a low amount of fluorescence background in the corral region. Using these relationships, the fluorescence intensity and the fluorescence lifetime in FLIM images were used to calculate the fluorophore concentration at each location in corrals before or after electrophoresis.

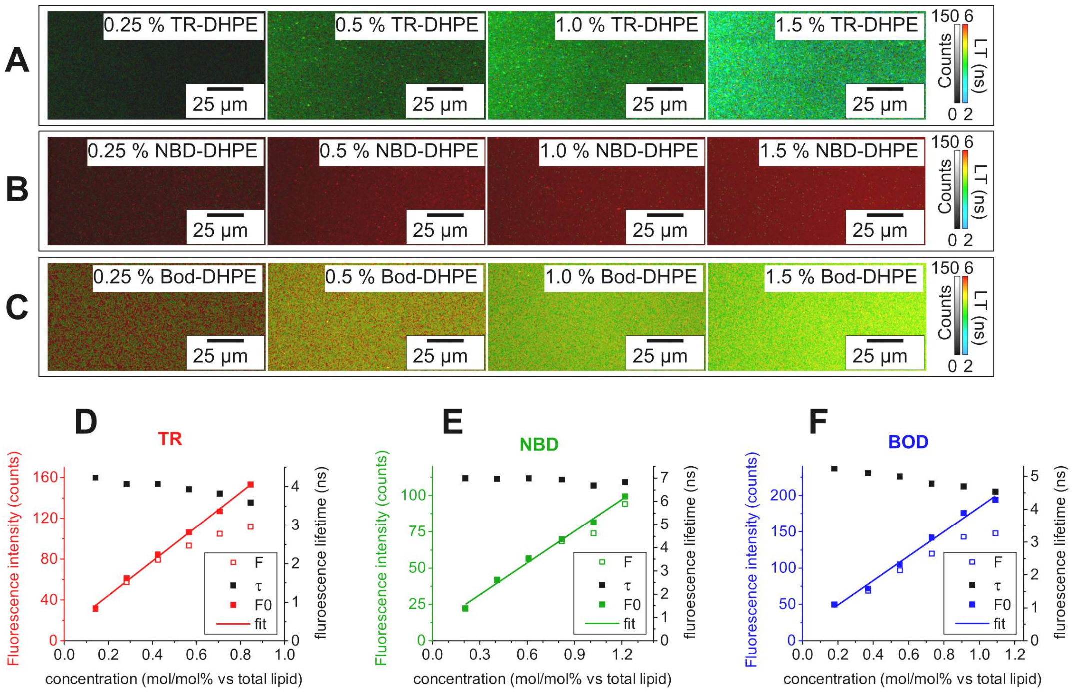


***Figure S3:*** *FLIM images of lipid bilayer standard samples containing fluorophores at defined concentrations used to generate standard curves from which to estimate fluorophore concentrations.* ***(A)*** *Example FLIM images of a series of SLBs containing increasing concentrations of TR-DHPE (concentrations in weight/weight, as labelled). FLIM images were obtained using the same settings as for in-membrane electrophoresis experiments.* ***(B)*** *Example FLIM images of a series of lipid bilayers containing increasing concentrations of NBD-DHPE.* ***(C)*** *Example FLIM images of a series of lipid bilayers containing increasing concentrations of BODIPY-DHPE.* ***(D)-(F)*** *The raw fluorescence intensity (F) observed within the sample (counts/pix) plotted against the known fluorophore concentration of the SLB. Corrected "non-quenched" fluorescence intensity ($F_0$) was calculated from F using main text Eq. 22. $\tau$ was plotted as measured from the FLIM data. All numerical data for F, $F_0$ and $\tau$ is in* ***Table S3****.*

| Sample | Conc. (% wt/wt) | Conc. (%mol/mol) | *F* (counts /pix) | $\tau$ (ns) | *F0* (counts/pix) |
|---|---|---|---|---|---|
| TR | 0.25 | 0.14 | 31.7 | 4.23 | 31.3 |
| | 0.50 | 0.28 | 57.5 | 4.07 | 61.2 |
| | 0.75 | 0.42 | 79.4 | 4.07 | 84.6 |
| | 1.00 | 0.57 | 93.1 | 3.93 | 106.3 |
| | 1.25 | 0.71 | 105.0 | 3.82 | 126.9 |
| | 1.50 | 0.85 | 112.0 | 3.59 | 153.3 |
| NBD | 0.25 | 0.21 | 22.1 | 6.99 | 22.2 |
| | 0.50 | 0.41 | 41.6 | 6.97 | 42.0 |
| | 0.75 | 0.61 | 56.3 | 6.98 | 56.6 |
| | 1.00 | 0.82 | 68.4 | 6.93 | 69.8 |
| | 1.25 | 1.02 | 74.0 | 6.67 | 81.5 |
| | 1.50 | 1.22 | 94.3 | 6.82 | 99.3 |
| BODIPY | 0.25 | 0.18 | 50.0 | 5.23 | 49.4 |
| | 0.50 | 0.37 | 68.8 | 5.09 | 71.8 |
| | 0.75 | 0.55 | 96.8 | 4.99 | 105.1 |
| | 1.00 | 0.73 | 120.0 | 4.78 | 142.0 |
| | 1.25 | 0.91 | 143.0 | 4.69 | 175.8 |
| | 1.50 | 1.09 | 148.0 | 4.54 | 194.2 |

***Table S3 (caption on next page)***

***Table S3:*** *Calculation of the "non-quenched" fluorescence intensity as a function of fluorophore concentration for TR, NBD and BODIPY. The fluorophore concentration in weight/weight was converted to mole/mole using the molecular weights of each lipid, DOPC, TR-DHPE, NBD-DHPE and BODIPY-DHPE (786, 1382, 956 and 1067 g/mol, respectively). The raw fluorescence intensity, F, and the fluorescence lifetime, $\tau$, were measured for each image (average of all pixels in one 25-frame FLIM acquisition). The non-quenched intensity, $F_0$, was calculated from F and $\tau$ using the equation:* $F_0 = F \cdot \exp\left[2\ln\left(\frac{\tau_0}{\tau}\right)\right]$.

### SI 4. Generation of concentration profiles and quantifying quenching from FLIM images of lipid bilayers undergoing electrophoresis

To generate a concentration profile of the fluorophore distribution across a membrane corral, the conversion factors specified in the previous section (**SI 3**) were applied. Below, an example of the conversion process is described for TR (the data and related text is adapted from ref. [3]). This process was applied to all samples (TR, NBD, BODIPY, new data). **Figure S4A** shows a FLIM image of a membrane corral at electrophoretic equilibrium which had a starting concentration of 0.28% (mol/mol) TR, showing the direction of the profiles generated from left-to-right across the membrane (*white dashed region*). The mean values for fluorescence intensity and lifetime were calculated for each horizontal (x) position across the membrane corral by averaging vertical columns (y) of pixels. The resulting profiles for fluorescence lifetime and raw fluorescence intensity are shown in **Figure S4B-C** (*black* and *bright red* datapoints, respectively). The corrected fluorescence intensity ($F_0$) profile was calculated using main text *Eq. 22*, as described in the previous SI section (*dark red datapoints* in **Figure S4C**). The mole-to-mole TR fluorophore concentration at each horizontal position (*blue datapoints* in **Figure S4D**) was then calculated from $F_0$, as above. We found that the TR concentration in this example increased roughly exponentially from zero on the right side of the corral up to ~2.0% (mol/mol) at the left side of the corral. This was correlated to a decrease in the fluorescence lifetime of TR from ~4 ns to ~2.8 ns from right-to-left.

To compare the fluorescence quenching between TR, NBD and BODIPY this analysis was performed for all the membrane corral samples described in the main text. The profiles of raw fluorescence intensity and fluorescence lifetime are shown in **Figure S4E** and **Figure S4F**. As expected, all fluorophores show much higher fluorescence intensity at the left-edge of the corral correlated to a lowered fluorescence lifetime. The corrected fluorescence intensity and the calculation concentration profiles (**Figure S4G-H**). The maximum concentrations achieved at the left edge of the corral vary somewhat from 2-3% for TR and BODIPY to 4-6% for NBD due to different mole-to-mole starting concentrations for the different fluorophores. The important point is not the absolute concentration achieved but what degree of quenching is observed for that concentration. The amount of quenching can be quantified from the fluorescence lifetimes as the Quenching Efficiency ($QE = 1 - \tau/\tau_0$) and profiles were plotted of $QE$ against distance (**Figure S4I**). A side-by-side comparison of the concentration profiles versus the $QE$ profiles is instructive (**Figure S4**, panel **H** *vs.* **I**). It is clear that BODIPY becomes most quenched for its concentration compared to NBD which has much lower quenching despite reaching double the concentration; TR appears to have intermediate levels of quenching.

Whilst comparison of concentration profiles versus *QE* profiles is revealing, we decided that that it would be even more informative to directly compare the fluorophore concentration to the *QE* on the same graph. This would allow the concentration dependence of quenching to be assessed and related to the theoretical models, so that one could assess whether the transfer-to-trap mechanism is occurring. Therefore, scatter plots of $QE$ against concentration were generated by using the fact that the line profiles described above are actually paired data, i.e., each location on a concentration profile has an observed value for fluorescence lifetime. Plots of $QE$ against concentration are displayed in the main text as part of **Figure 4** and **Figure 5** and are explained and discussed there.

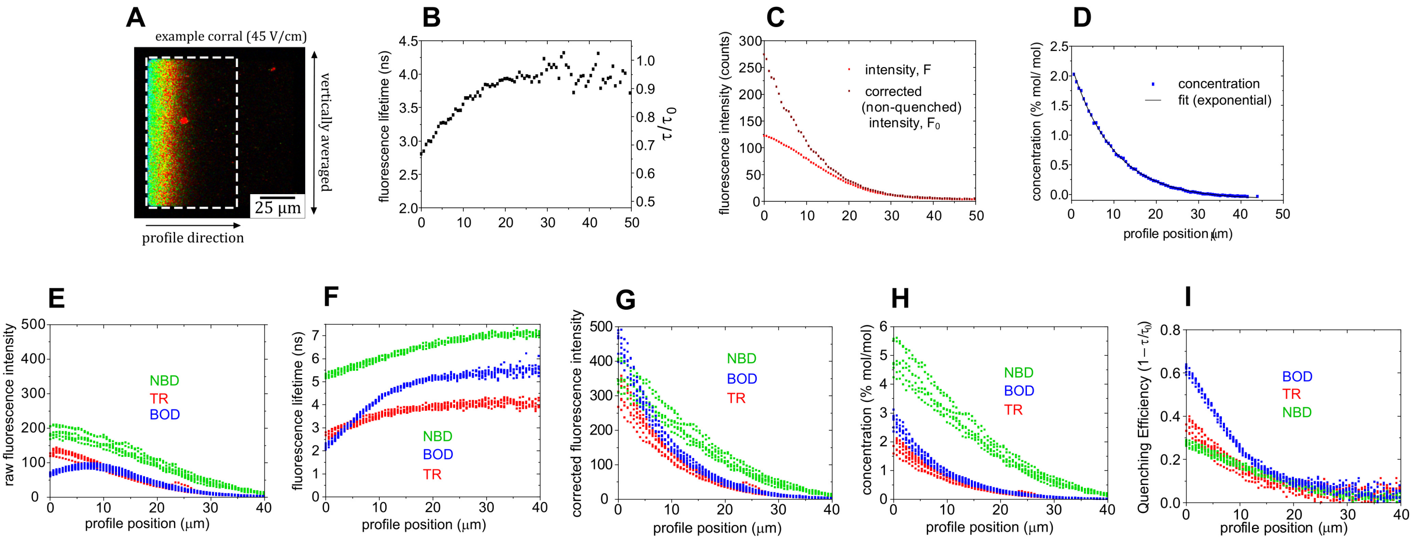


***Figure S4:*** *Method for generating concentration profiles from a FLIM image applied to TR, NBD and BODIPY.* ***(A)*** *Example FLIM image of a 0.28% (mol/mol) TR corral at equilibrium in an E-field. The white dashed box denotes the region-of-interest (ROI) from which horizontal profiles of lifetime and intensity were obtained by averaging the pixels accumulated vertically (typically 150 pix/vertical, improving the signal-to-noise).* ***(B)*** *Profile of fluorescence lifetime against x-position obtained from the white dashed region in (A).* ***(C)*** *Profile of raw fluorescence intensity profile (light red), F, obtained from the white dashed region in (A). The corrected fluorescence intensity, $F_0$ (dark red), was calculated from the profile for $F(x)$ and the profile for $\tau(x)$ using a Eq. 22 in the following form: $F_0(x) = F(x) \cdot e^{2\,ln\,[\tau_0/\tau(x)]}$.* ***(D)*** *The concentration profile (blue) calculated from the data for $F_0$ from (C) using the direct proportionality relationship between the molar concentration and non-quenched intensity of a fluorophore, $C = F_0 / 168.3$. The solid black line is a fit to mono-exponential function: $C(x) = a \cdot e^{-bx} + y_0$ (a, b and $y_0$ are fitting constants).* ***(E)*** *The raw fluorescence intensity profiles from a series of membrane corrals containing either TR, NBD or BODIPY fluorophores (colouring as labelled). Each profile is generated by vertically averaging the data, as shown in the example membrane corral in (A). Data from multiple independent membrane corrals is overlaid (n = 6 for each fluorophore type).* ***(F)*** *Fluorescence lifetime profiles corresponding to the fluorescence intensity profiles from (E).* ***(G)*** *Profiles of corrected fluorescence intensity as calculated from the profiles for $F(x)$ from (E) and the profiles for $\tau(x)$ from (F). Before calculating these profiles, a few counts of noise were subtracted from the raw fluorescence intensity by aligning all curves to F = 0 at x > 40 µm.* ***(H)*** *Concentration profiles as calculated from the data for $F_0$ from (G) using the direct proportionality relationships between the molar concentration and corrected intensity of a fluorophore, $C = F_0 / 168.3$, $F_0 / 73.0$, or $F_0 / 168.8$ for TR, NBD or BODIPY, respectively.* ***(I)*** *Profiles of Quenching Efficiency calculating from the equation: $QE(x) = 1 - [\tau(x)/\tau_0]$ using the corresponding fluorescence lifetime data from (F). Panels A-D are reproduced from Meredith et al. 2023 [3]. All other data is new.*

### SI 5. Comparison of the simplified versus exact mathematical models for quenching

To recap, the reduction in fluorescence intensity and lifetimes according to main text *Eq. 5* and *Eq.7*:

$$\frac{\tau}{\tau_0} = 1 - P_{ETT} \qquad \text{main text } Eq.\ 5$$

and:

$$\frac{F}{F_0} = (1 - P_{ETT})(1 - f_T) \qquad \text{main text } Eq.\ 7$$

where $f_T$ is the fraction of fluorophores involved in traps, $P_{ETT}$ is the probability of excitation transfer to a trap site, $F_0$ is the original fluorescence intensity, $F$ is the fluorescence intensity after quenching, $\tau_0$ is the original fluorescence lifetime, $\tau$ is the fluorescence lifetime after quenching.

$f_T$ depends only on the critical radius of trap formation ($R_C$) for a given fluorophore, at a given concentration ($C_N$). Whereas, $P_{ETT}$ depends upon the FRET efficiency which, in turn, depends upon the Förster radius ($R_0$).

**The exact model** applies an expression for $P_{ETT}$ based on the fluorophore concentration ($C_N$), the known value for Förster radius ($R_0$), and the critical radius of trap formation ($R_C$), so:

$$\frac{\tau}{\tau_0} = 1 - \left[\frac{R_0^6}{R_0^6 + (\pi C_N)^{-3}}\left(1 - e^{-\pi R_C^2 C_N}\right)\right] \qquad \text{Eq. S1}$$

and:

$$\frac{F}{F_0} = \left\{1 - \left[\frac{R_0^6}{R_0^6 + (\pi C_N)^{-3}}\left(1 - e^{-\pi R_C^2 C_N}\right)\right]\right\}\left\{e^{-\pi R_C^2 C_N}\right\} \qquad \text{Eq. S2}$$

Whereas, **the simplified model** assumes that $P_{FRET} = 1$, therefore $P_{ETT} = f_T$, and so:

$$\frac{\tau}{\tau_0} = e^{-\pi R_C^2 C_N} \qquad \text{main text Eq. 16}$$

and:

$$\frac{F}{F_0} = e^{-2\pi R_C^2 C_N} \qquad \text{main text Eq. 20}$$

At the end of this study, values for the critical radius for trap formation ($R_C$) were calculated from the experimental data of FLIM analysis of quenching during fluorophore electrophoresis (see main text **Figure 5**). These $R_C$ values can be used to simulate the reduction in fluorescence intensity and the reduction in fluorescence lifetime that would occur according to the transfer to trap and statistical pair model, by solving both the exact models (Eq. S1 and Eq. S2) and the simplified model (Eq. 16 and Eq. 20) for a range of concentrations ($C_N$). The results are shown in **Figure S5**, below.

Here, we see the relative fluorescence intensity ($F/F_0$) decreases more rapidly with concentration than the relative fluorescence lifetime ($\tau/\tau_0$) and generally the trend lines for NBD > TR > BODIPY (**Figure S5** vs **5B**). There is a slight deviation between the trend produced by the exact model as compared to the simplified model at the lower concentrations, where the FRET assumption is invalid, and this deviation is most pronounced for NBD because it has the lowest Förster radius. Overall, these discrepancies are minor and equate to differences of 10% in the worst cases and the agreement between the two models is generally very good.

One subtle difference worth noting is that, according to the exact model, the fluorescence lifetime does not change appreciably below a fluorophore concentration of ~0.75% whereas the fluorescence intensity becomes somewhat quenched immediately. Now that we have estimates for $R_0$ and $R_C$ for all three fluorophores, these graphs (solid lines in **Figure S5**) may be considered the most accurate representation of the expectations for fluorescence from these fluorophores.

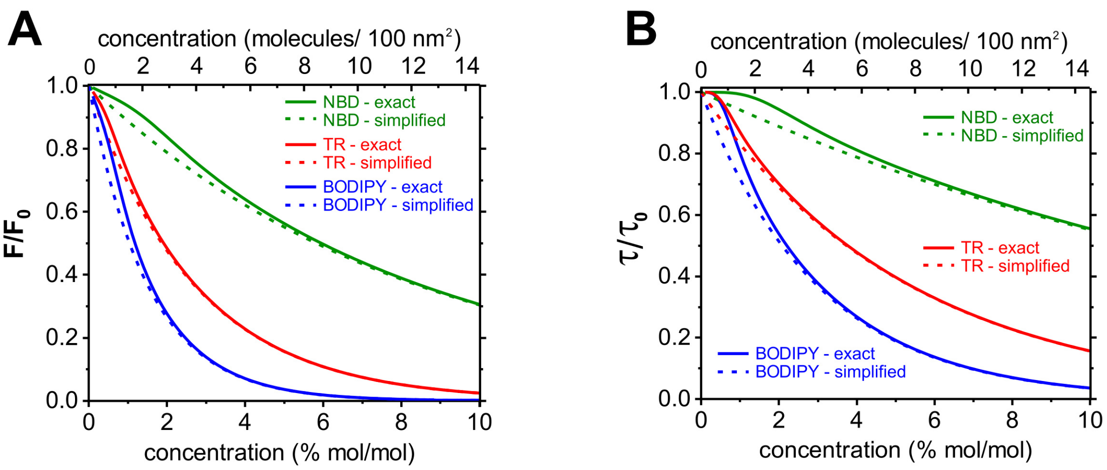


***Figure S5:*** *Comparison of the exact and simplified theoretical models for transfer-to-trap quenching.* ***(A)*** *Relative fluorescence intensity as calculated from either Eq. S2 (exact) or Eq. 20 (simplified). This is proportional to the fluorescence quantum yield.* ***(B)*** *Relative fluorescence lifetime as calculated from either Eq. S1 (exact) or Eq. 16 (simplified). "Relative fluorescence lifetime" is sometimes termed the "fluorescence lifetime yield".*
